# An unusual activity of mycobacterial MutT1 Nudix hydrolase domain as a protein phosphatase regulates nucleoside diphosphate kinase (NDK) function

**DOI:** 10.1101/2024.09.26.615297

**Authors:** Elhassan Ali Fathi Emam, Koyel Roy, Devendra Pratap Singh, Deepak K Saini, Umesh Varshney

## Abstract

MutT proteins are Nudix hydrolases characterized by the presence of a Nudix box, GX5EX7REUXEEXGU, where U is a bulky hydrophobic residue and X is any residue. Major MutT proteins hydrolyse 8-oxo-(d)GTP (8-oxo-GTP or 8-oxo-dGTP) to the corresponding 8-oxo-(d)GMP, preventing their incorporation into nucleic acids. Mycobacterial MutT1 comprises an N-terminal domain (NTD) harbouring the Nudix box motif, and a C-terminal domain (CTD) harbouring the RHG histidine phosphatase motif. Interestingly, unlike other MutTs, the MutT1 hydrolyses the mutagenic 8-oxo-(d)GTP to corresponding 8-oxo-(d)GDP. Nucleoside diphosphate kinase (NDK), a conserved protein, carries out reversible conversion of (d)NDPs to (d)NTPs through phospho-NDK (NDK-*Pi*) intermediate. Recently, we showed that NDK-*Pi* converts 8-oxo-dGDP to 8-oxo-dGTP and escalates A to C mutations in a MutT deficient *Escherichia coli*. We now show that both *Mycobacterium tuberculosis* MutT1, and *M. smegmatis* MutT1, through their NTD (Nudix hydrolase motifs) function as protein phosphatase to regulate the levels of NDK-*Pi* to NDK and prevent it from catalysing conversion of (d)NDPs to (d)NTPs (including conversion of 8-oxo-dGDP to 8-oxo-dGTP). To corroborate this function, we show that *Msm*MutT1 decreases A to C mutations in *E. coli* under the conditions of *Eco*NDK overexpression.

**Importance:** MutT proteins, having a Nudix box domain, hydrolyse the mutagenic 8-oxo-dGTP to 8-oxo-dGMP. However, mycobacterial MutT (MutT1) comprises an N-terminal domain (NTD) harboring a Nudix box, and a C-terminal domain (CTD) harboring an RHG histidine phosphatase. Unlike other MutTs, mycobaterial MutT1 hydrolyses 8-oxo-dGTP to 8-oxo-dGDP. Nucleoside diphosphate kinase (NDK), a conserved protein, converts 8-oxo-dGDP to 8-oxo-dGTP through phospho-NDK (NDK-*Pi*) intermediate, and escalates A to C mutations. Here, we show that the mycobacterial MutT1 is unprecedented in that its NTD (Nudix box), functions as protein phosphatase to regulate NDK-*Pi* levels and prevents it from converting dNDPs to dNTPs (including 8-oxo-dGDP to 8-oxo-dGTP conversion). In addition, mycobacterial MutT1 decreases A to C mutations in *Escherichia coli* under the conditions of NDK overexpression.

## Introduction

*Mycobacterium tuberculosis*, a pathogenic bacterium, homes the host macrophages, where it encounters high levels of reactive oxygen species (ROS) and reactive nitrogen intermediates (RNI) generated as part of the host innate immune response [1, 2]. ROS and RNI can damage nucleotide bases in nucleic acids and the free nucleotide pool. Due to its low redox potential, guanine (G) is highly susceptible to oxidative damage. Oxidation of (d)GDP (GDP/dGDP) or (d)GTP (GTP/dGTP) converts them to 8-oxo-(d)GDP or 8-oxo-(d)GTP, respectively [3–5]. As 8-oxo-dGTP can pair with A, its misincorporation into the genome may result in AT to CG transversion mutations [6, 7].

The 8-oxo-(d)GDP can also be converted to 8-oxo-dGTP by nucleoside diphosphate kinase (NDK) [8]. NDK is an evolutionarily conserved enzyme responsible for multiple cellular functions. In addition, the pathogens like *M. tuberculosis*, also secrete NDK (*Mtb*NDK) as an effector protein that regulates diverse host cellular processes, including phagocytosis, apoptosis/necrosis, and inflammatory response [9, 10]. *Mtb*NDK forms a stable hexamer [11]. A major role of NDK is to maintain (homeostasis) the intracellular nucleoside diphosphate and nucleoside triphosphate pools [12–14]. The catalysis by NDK involves autophosphorylation of a conserved His in it by the γ-phosphate of nucleoside triphosphates. The phosphate from His-*Pi* is then transferred to nucleoside diphosphates through a ping-pong mechanism, converting them into nucleoside triphosphates [14, 15]. Recently, we observed that NDK catalyses formation of 8-oxo-dGTP from 8-oxo-dGDP at a rate 2-3 times higher than its activity in converting 8-oxo-dGTP into 8-oxo-dGDP [8]. However, no significant differences were observed in the rates of reversible conversions between GTP and GDP. Importantly, overexpression of NDK in *Escherichia coli* Δ*mutT* increases A to C mutations [8].

MutT proteins belong to the Nudix hydrolase family, which possess a Nudix hydrolase motif (GX5EX7REUXEEXGU). In this motif, U indicates a bulky hydrophobic residue, whereas X represents any residue [16, 17]. *E. coli* MutT hydrolyses 8-oxo-(d)GTP and 8-oxo-(d)GDP into 8-oxo-(d)GMP [18]. In mycobacteria, four of the Nudix hydrolases (MutT1, MutT2, MutT3 and MutT4) were identified as orthologs of *E. coli* MutT. Of these, MutT1 hydrolyses 8-oxo-(d)GTP into 8-oxo-(d)GDP. Hydrolysis of 8-oxo-dGTP to 8-oxo-dGDP also prevents its misincorporation into the genome [5, 6]. Mycobacterial MutT1 complements *E. coli* CC101 (Δ*mutT*) strain and decreases A to C mutations [19]. MutT2 functions majorly as an efficient dCTPase [14]. It has been reported that the mycobacterial MutT3 (RenU) is required for the pathogen to survive under the environment of oxidative stress inside macrophages [15]. The in vitro experiments revealed that MutT4 exhibits greater efficacy in the hydrolysis of adenine. Its deficiency in the *mutT4* mutant strain, results in a rise in the ratio of A to G (or T to C) mutations under oxidative stress [16, 17]. Unlike *E. coli* MutT or the mycobacterial MutT2, MutT3 and MutT4, the mycobacterial MutT1 is unusual in consisting of an N-terminal domain (NTD) harbouring a Nudix hydrolase motif, and a C-terminus domain (CTD) harbouring an RHG histidine phosphatase motif of unknown activity [20, 21].

Although NDK is involved in diverse biological processes (9) and is autophosphorylated (to NDK-*Pi*) on a conserved His, no studies are available that have explored its possible regulation by dephosphorylation of its phospho-His by any of the cellular proteins. Because of the cross-talk between MutT and NDK in *E. coli*, we wondered if NDK could be a substrate for dephosphorylation by mycobacterial MutT1.

Our biochemical and in vivo studies, designed to investigate the possible roles of the N-terminal domain (NTD), and C-terminal domain (CTD) of MutT1 in regulation of NDK, reveal that NDK-*Pi* is a substrate for dephosphorylation by MutT1. While the dephosphorylation of NDK-*Pi* is catalysed by NTD of MutT1, we show that the CTD of MutT1 is important in supporting the phosphatase activity of the NTD. Finally, we show how MutT1 regulates the activity of NDK both in vitro and in vivo.

## Results

### Mycobacterial MutT1 exhibits phosphatase activity against NDK-*Pi*

To test if autophosphorylated NDK (NDK-*Pi*) is a substrate for MutT1 phosphatase activity, we purified *Msm*NDK, *Mtb*NDK, *Msm*MutT1, and *Mtb*MutT1 (Fig. S1A). As reported earlier [19, 21], in this study too we observed that *Msm*MutT1 and *Mtb*MutT1 migrate as doublets on SDS-PAGE. The assays for the phosphatase activity using *p*NPP as a nonspecific substrate (Fig. S1B) revealed that both the MutT1 proteins were active (compare bars 2 and 3 with bar 1 control). Incubation of *Msm*NDK-*Pi* with *Msm*MutT1 showed that the MutT1 resulted in dephosphorylation of *Msm*NDK-*Pi* (Fig. 1A, *panel i*, compare lane 2 with lane 1). SDS-PAGE analysis of the same gel (Fig. 1A, *panel ii*), showed that equal amounts of *Msm*NDK were present in both the lanes, and that its dephosphorylation occurred only in the sample containing *Msm*MutT1. Quantification of the gel using phosphor imager showed that *Msm*NDK-*Pi* dephosphorylation was ∼90% (shown in the bottom of *panel i*; Fig. S9A). When we repeated the assay with *Mtb* proteins (*Mtb*NDK and *Mtb*MutT1), identical observations were made (Fig. 1B), revealing that MutT1 has a phosphatase activity against NDK-*Pi*.

**Figure 1:**
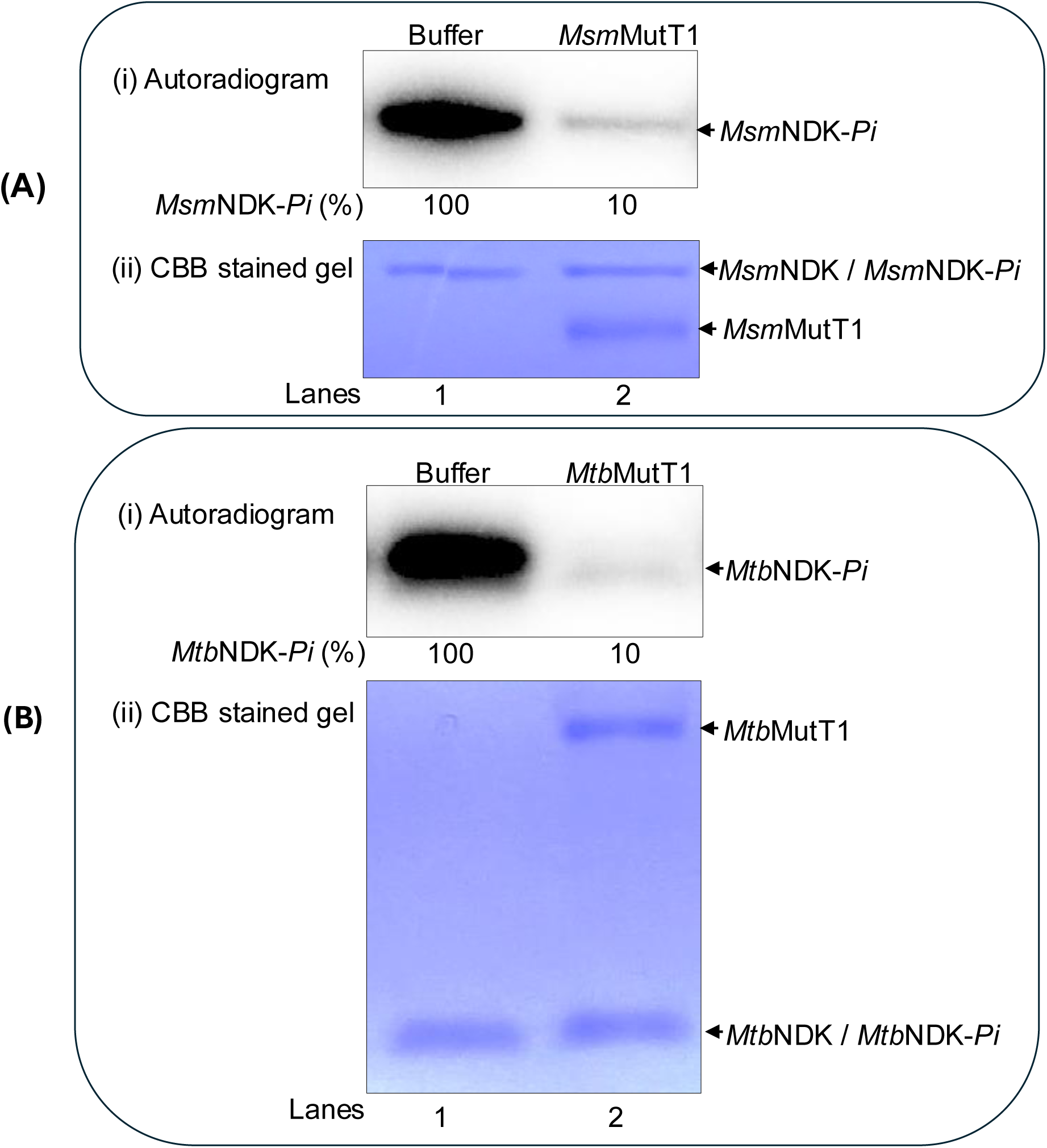
Dephosphorylation of NDK-*Pi* with mycobacterial MutT1. (A) Dephosphorylation of *Msm*NDK-*Pi* by *Msm*MutT1. *Msm*NDK was autophosphorylated in the presence of γ-^32^P-ATP to obtain *Msm*NDK-*Pi*. The *Msm*NDK-*Pi* (1μg) was then incubated with either buffer alone (lane 1) or 1μg *Msm*MutT1 (lane 2) at 30 °C for 1 h, mixed with 5 μL SDS-PAGE sample buffer and analysed on 12% SDS-PAGE (because the phospho-His is labile to heat, the samples were not heated). Gels were subjected to phosphor imaging, fixed, and stained with Coomassie brilliant blue (CBB). Panels (i) and (ii) represent autoradiogram, and CBB stained gel, respectively. Numbers (%) below *panel i* show the phosphorylation levels of *Msm*NDK-*Pi* (see also Fig. S9A). **(B) Dephosphorylation of *Mtb*NDK-*Pi* by *Mtb*MutT1.** The *Mtb*NDK-*Pi* (1 μg) obtained as in (A), was incubated with buffer alone (lane 1) or *Mtb*MutT1 (1μg) (lane 2). The reactions were processed as in (A) above. Panels: (i) autoradiogram, (ii) CBB stained gel. Numbers (%) below *panel i* show the phosphorylation levels of *Mtb*NDK-*Pi* (see also Fig. S9B).

### Generation of single site mutations in NTD and CTD of MutT1 proteins

Mycobacterial MutT1comprises two domains, NTD having Nudix hydrolase motif, and CTD having RHG histidine phosphatase motif. To better understand the role of MutT1 in dephosphorylation of NDK-*Pi*, we generated single site mutations of E81A (in NTD) or H170A (in CTD) in *Msm*MutT1 (Fig S2A). The E81 and H170 are critical residues in the Nudix box and the RHG His-phosphatase motifs, respectively [22–24]. The assays using the mutant proteins (Fig. S2B) showed that the E81A mutation in NTD rendered MutT1 inactive in dephosphorylation of *p*NPP (Fig. S2C, compare bar 3 with bars 1 and 2). However, the H170A mutation in CTD had no impact on the dephosphorylation of *p*NPP (Fig. S2C, compare bar 4 with bars 1 and 2). The equivalent mutations in *Mtb*MutT1 corresponded to E69A in Nudix hydrolase motif in NTD, and H161A in RHG motif in CTD (Fig. S3A). Consistent with the results on *Msm*MutT1 mutants, *Mtb*MutT1 E69A but not the *Mtb*MutT1 H161A lost its dephosphorylation activity on *p*NPP (Figs. S3B and S3C).

### Nudix hydrolase motif is the source of protein phosphatase activity on NDK-*Pi*

To check which of the domains of MutT1 exhibited the phosphatase activity on NDK-*Pi*, *Msm*NDK-*Pi* was treated with *Msm*MutT1, *Msm*MutT1(E81A) or *Msm*MutT1(H170A). The results in Fig. 2A (*panels i* and *ii*) show that dephosphorylation of *Msm*NDK-*Pi* occurred upon treatment with *Msm*MutT1 and *Msm*MutT1 H170A (Fig. 2A, *panel i*, lanes 2 and 4), respectively. No significant dephosphorylation was detected when the *Msm*NDK-*Pi* was incubated with *Msm*MutT1 E81A (Fig. 2A, *panel i*, compare lane 3 with the buffer alone lane 1; see also Fig. S9C). Similar results were obtained when *Mtb*NDK-*Pi* was treated with *Mtb*MutT1, *Mtb*MutT1 E69A or *Mtb*MutT1 H161A (Fig. 2B, *panel i*). The SDS-PAGE analysis (Fig. 2A and 2B, *panels ii*) shows that equal amounts of NDK and MutT1 proteins were used. Taken together, these findings show that mutation in NTD Nudix box (*Msm*MutT1 E81A and *Mtb*MutT1 E69A) resulted in loss of their dephosphorylation activities on NDK-*Pi* suggesting that the Nudix hydrolase motif of the MutT1 proteins is responsible for dephosphorylation of NDK-*Pi*. *Msm*NDK and *Mtb*NDK share ∼80% sequence similarity (Fig. S4A). Not unexpectedly, we observed efficient dephosphorylation of *Msm*NDK-*Pi* by both *Msm*MutT1, *Mtb*MutT1 and their CTD mutants but not the NTD mutants (Fig. S4B). Likewise, *Mtb*NDK-*Pi* is dephosphorylated with either of the MutT1 or their CTD mutants but not the NTD mutants (Fig. S4C). These data show that MutT1 proteins act not only on their homologous NDKs but also on those from different mycobacterial species. Importantly, these observations further consolidate the role of MutT1 NTD in NDK-*Pi* dephosphorylation.

**Figure 2:**
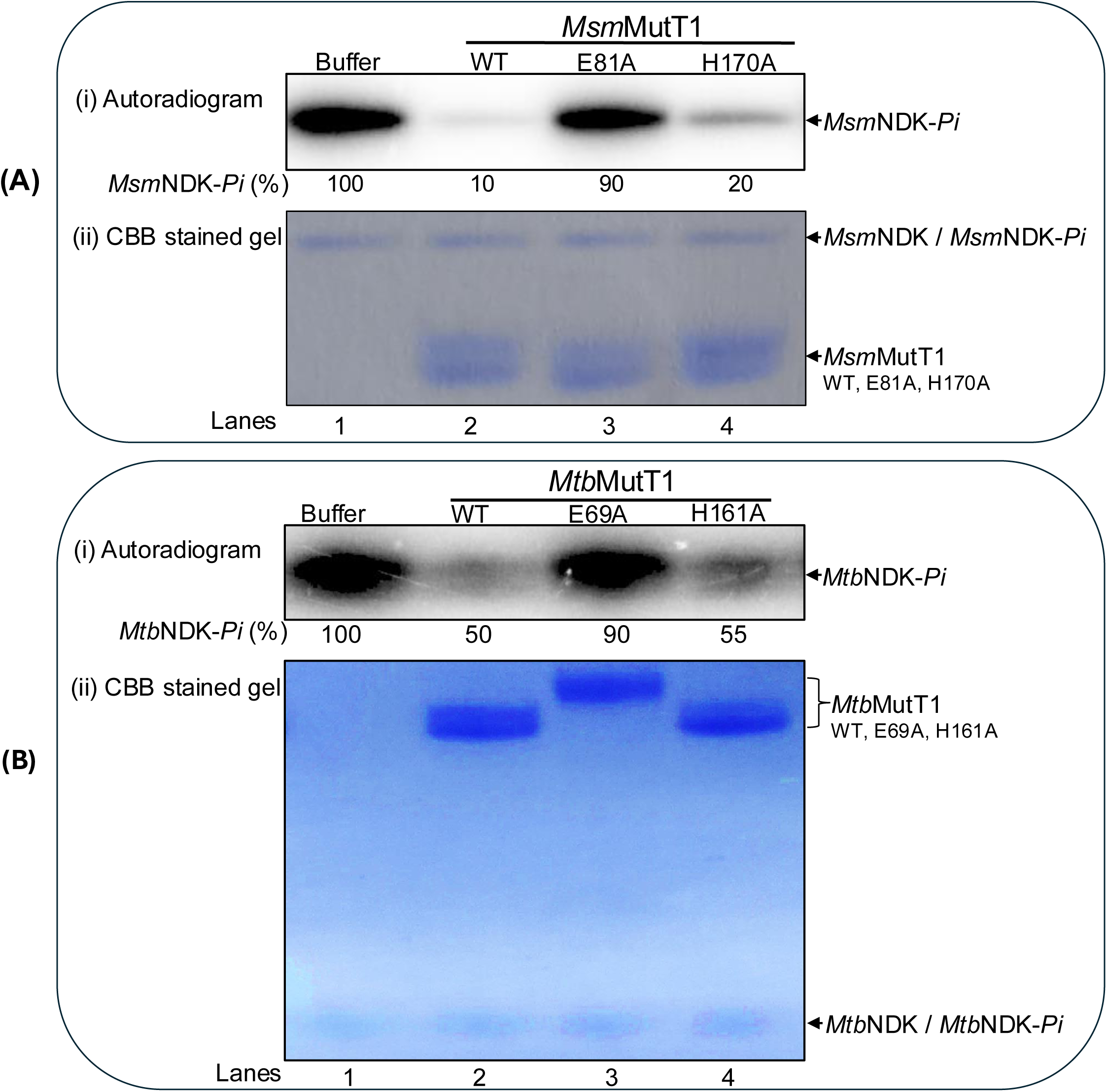
Dephosphorylation of NDK-*Pi* with MutT1 mutants. **(A) Dephosphorylation of *Msm*NDK-*Pi* by *Msm*MutT1 (WT, E81A, and H170A).** The autophosphorylated *Msm*NDK-*Pi* (1 μg) was incubated with buffer alone (lane 1) or 2 μg *Msm*MutT1 (lane 2), or 2 μg *Msm*MutT1 E81A (lane 3), or 2 μg *Msm*MutT1 (H170A) (lane 4). Panels (i) and (ii) represent autoradiogram and CBB stained gel. NDK-*Pi* (%) levels are shown below *panel i*. For further quantification of *Msm*NDK-*Pi* levels refer to Fig. S9C. **(B) Dephosphorylation of *Mtb*NDK-*Pi* by *Mtb*MutT1 (WT, E69A, and H161A).** The *Mtb*NDK-*Pi* (1 μg) was incubated with buffer alone (lane 1), or 2 μg *Mtb*MutT1 (lane 2), or 2 μg *Mtb*MutT1 E69A (lane 3), or 2 μg *Mtb*MutT1 H161A (lane 4). Panels (i) and (ii) represent autoradiogram, and CBB stained gel, respectively. NDK-*Pi* (%) levels are shown below *panel i*. For further quantification of *Msm*NDK-*Pi* levels refer to Fig. S9D. Three technical replicates were used in this experiment. It may be noted that in lane 1, because the samples are not heated after adding sample dye, *Msm*NDK migrates much slowly than its expected monomeric molecular weight (also refer to Fig. S8). Lane 2: *Msm*MutT1E81A migrates slightly faster than *Msm*MutT1, while *Mtb*MutT1 migrates slower than *Mtb*MutT1. This was only observed when samples are not heated in the sample buffer.

### MutT1 CTD supports the phosphatase activity of MutT1 NTD

The observations (Fig. 2) that the Nudix hydrolase is responsible for dephosphorylation of NDK-*Pi* raises a question if other MutT proteins (possessing a single domain corresponding to the mycobacterial MutT1 NTD) would act on NDK-*Pi*. Thus, we purified *Eco*MutT, *Msm*MutT2, and *Eco*NDK (Fig. S5A). Both *Eco*MutT and *Msm*MutT2 showed a strong phosphatase activity on *p*NPP (Fig. S5B). However, no dephosphorylation of *Eco*NDK-*Pi* was detected when treated with *Eco*MutT (Fig. 3A, *panels* i-ii). Similarly, no dephosphorylation of *Msm*NDK-*Pi* occurred with *Msm*MutT2 (Fig. 3B, *panels* i-ii). To better understand why *Eco*MutT may not dephosphorylate *Eco*NDK-*Pi*, we made use of AlphaFold Colab protein-protein interaction tool [25, 26]. We observed a single site interaction between *Eco*MutT and *Eco*NDK (Fig. S6A). While many interactions were observed between *Msm*NDK and *Msm*MutT1 (NTD) and (CTD) (Fig. S6B) indicating a possible requirement of MutT1 CTD in facilitating binding of NDK-*Pi* to MutT1 for dephosphorylation by MutT1 NTD.

**Figure 3:**
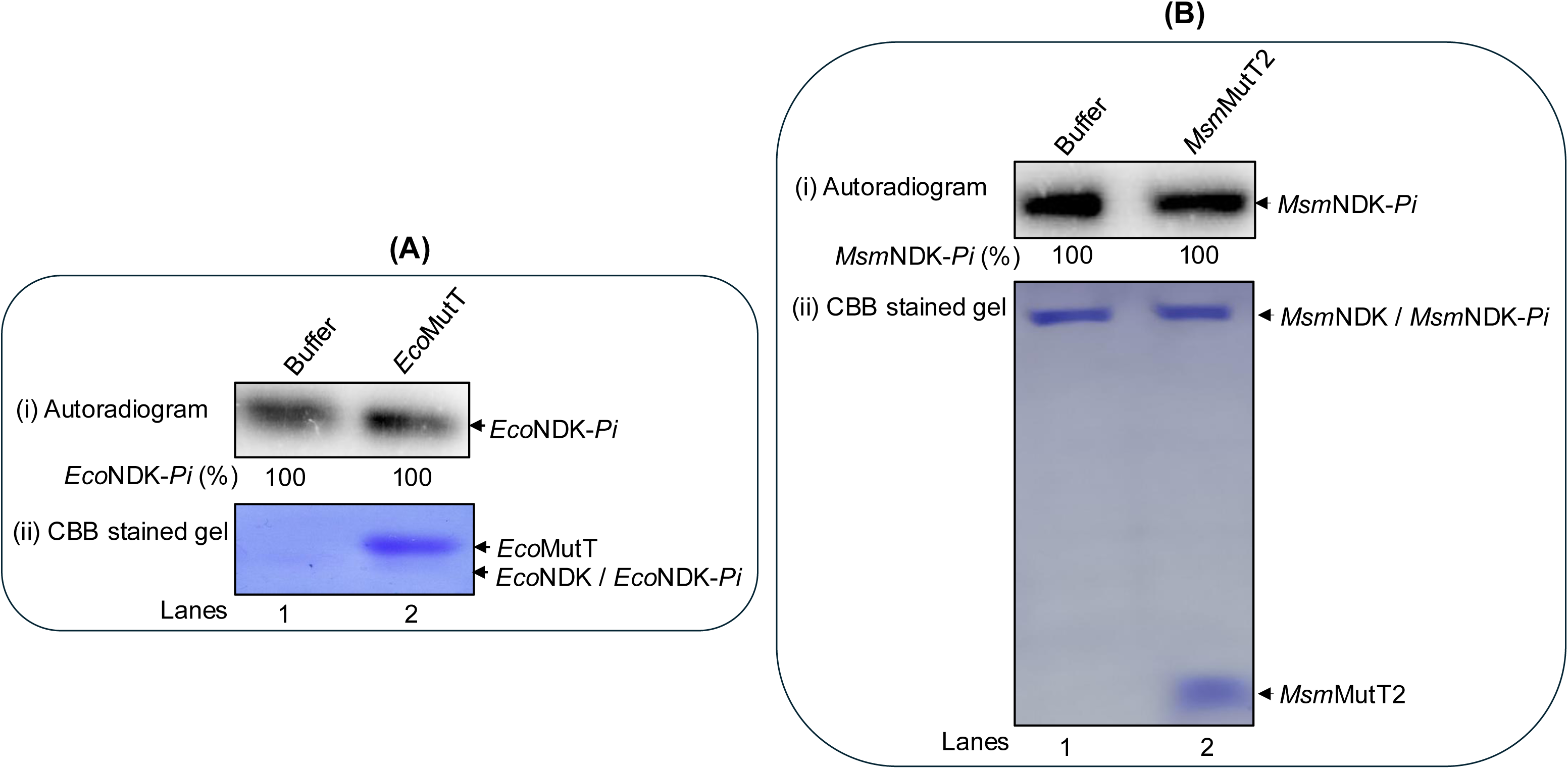
Treatments of NDK-*Pi* with *Eco*MutT and *Msm*MutT2. **(A) Treatment of *Eco*NDK-*Pi* with *Eco*MutT.** The *Eco*NDK-*Pi* (1 μg) was incubated with buffer alone (lane 1) or *Eco*MutT (2 μg) (lane 2). Panels (i) and (ii) represent autoradiogram and CBB stained gel. NDK-*Pi* (%) levels are shown below *panel i*. For further quantification of *Eco*NDK-*Pi* levels refer to Fig. S10A. **(B) Treatment of *Msm*NDK-*Pi* by *Msm*MutT2.** The *Msm*NDK-*Pi* (1 μg) was incubated with buffer alone (lane 1) or 1 μg *Msm*MutT2 (lane 2). Panels (i) and (ii) represent autoradiogram and CBB stained gel. *Msm*NDK-*Pi* (%) levels are shown below *panel i*. For further quantification of *Msm*NDK-*Pi* levels refer to Fig. S10B. It may be noted that because the samples are not heated in the sample buffer prior to loading in the SDS-PAGE, *Msm*NDK migrates slower than its expected monomeric molecular weight (also refer to Fig. S8).

To test this prediction, we generated a chimeric protein where NTD of *Msm*MutT1 was replaced with *Eco*MutT sequence. The chimera was purified and assayed for its general phosphatase activity using *p*NPP (Figs. S7A-S7D). All the MutT proteins (*Eco*MutT, *Msm*MutT1 and *Eco-Msm*MutT1 chimera) showed the general phosphatase activity (Fig. S7D, compare bars 2-4 with 1). However, *Eco*MutT showed the best activity and given that the Nudix hydrolase (NTD) conferred the phosphatase activity, the *Eco-Msm*MutT1, not unexpectedly, also showed a better activity compared with the *Msm*MutT1 (Fig. S7D, compare bars 3 and 4). Subsequently, *Eco*NDK-*Pi* was treated with *Eco*MutT, *Eco-Msm*MutT1 chimera, or *Msm*MutT1 (Fig. 4A). The results revealed a prominent decrease in the level of *Eco*NDK-*Pi* when incubated with *Eco-Msm*MutT1 chimera or *Msm*MutT1 (Fig. 4A. *panel i*, lanes 3 and 4). As control, no decrease in *Eco*NDK-*Pi* levels was detected upon its incubation with buffer alone or *Eco*MutT (Fig. 4A, *panel i*, lanes 1 and 2). Similar results were obtained when *Msm*NDK-*Pi* was used (Fig. 4B, *panels* i and ii). The observations strongly suggest that *Msm*MutT1 CTD structurally contributes to MutT1 NTD phosphatase activity on NDK-*Pi*.

**Figure 4:**
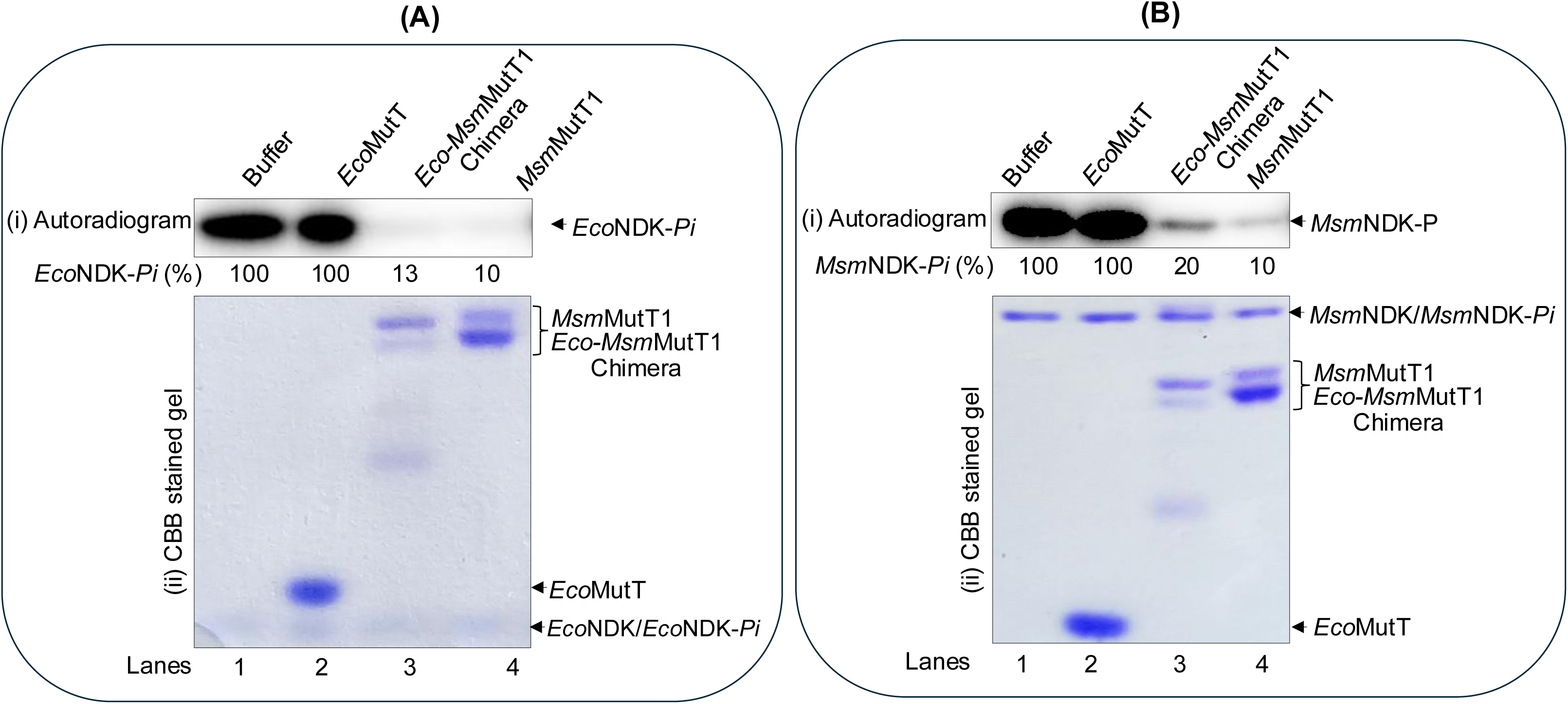
Dephosphorylation of NDK-*Pi* with *Eco-Msm*MutT1 chimera (A) Dephosphorylation of *Eco*NDK-*Pi* by *Eco*MutT, *Eco-Msm*MutT1 chimera and *Msm*MutT1. The autophosphorylated *Eco*NDK-*Pi* (0.5 μg) was incubated with buffer alone (lane 1), 2 μg *Eco*MutT (lane 2), 2 μg *Eco-Msm*MutT1 chimera (lane 3), or 2 μg *Msm*MutT1 (lane 4). Panels (i) and (ii) represent autoradiogram and CBB stained gel, respectively. NDK-*Pi* (%) levels are shown below *panel i*. For further quantification of *Eco*NDK-*Pi* levels refer to Fig. S10C. **(B) Dephosphorylation of *Msm*NDK-*Pi* by *Eco*MutT, *Eco-Msm*MutT1 chimera and *Msm*MutT1.** The *Msm*NDK-*Pi* (1 μg) was incubated with buffer alone (lane 1), 2 μg *Eco*MutT (lane 2), 2 μg *Eco-Msm*MutT1 chimera (lane 3), or 2 μg *Msm*MutT1 (lane 4). Panels (i) and (ii) represent autoradiogram and CBB stained gel. NDK-*Pi* (%) levels are shown below *panel i*. For further quantification of *Msm*NDK-*Pi* levels refer to Fig. S10D. Three technical replicates were used in this experiment. It may be noted that because the samples were not heated in the sample loading dye, *Msm*NDK migrates slower than its expected monomeric molecular weight (also refer to Fig. S8).

### MutT1 regulates NDK activity and its downstream function *in vitro*

NDK plays an important role in homeostasis of the cellular nucleotide pool crucial in maintaining genome integrity [8]. It executes reversible conversions of NDPs to NTPs. To understand the effect of MutT1 on this function of NDK, *Msm*NDK-*Pi* was incubated with *Msm*MutT1, *Msm*MutT1 E81A, and the *Eco-Msm*MutT1 chimera, and followed further by incubation with ADP. The results in Fig. 5A show that *Msm*NDK-*Pi* converted ADP to ATP when it was pre-incubated with buffer alone or the Nudix hydrolase dead mutant *Msm*MutT1 E81A (Fig. 5A, lanes 1 and 3). However, the activity of *Msm*NDK-*Pi* in converting ADP to ATP was highly suppressed upon its pre-treatment with *Msm*MutT1 or the *Eco-Msm*MutT1 chimera (Fig. 5A, lanes 2 and 4, respectively). The same results were obtained, when *Mtb*NDK-*Pi* was treated with the corresponding *Mtb*MutT1 proteins (Fig. 5B). We then asked if MutT1 restricts NDK-*Pi* from converting 8-oxo-dGDP to 8-oxo-dGTP. In this experiment also, *Mtb*MutT1 but not its E69A mutant, prevented conversion of 8-oxo-dGDP to 8-oxo-dGTP (Fig. 5C, compare lanes 2 and 3 with 1). These observations suggest that MutT1 regulates NDK activity in converting NDPs to NTPs.

**Figure. 5:**
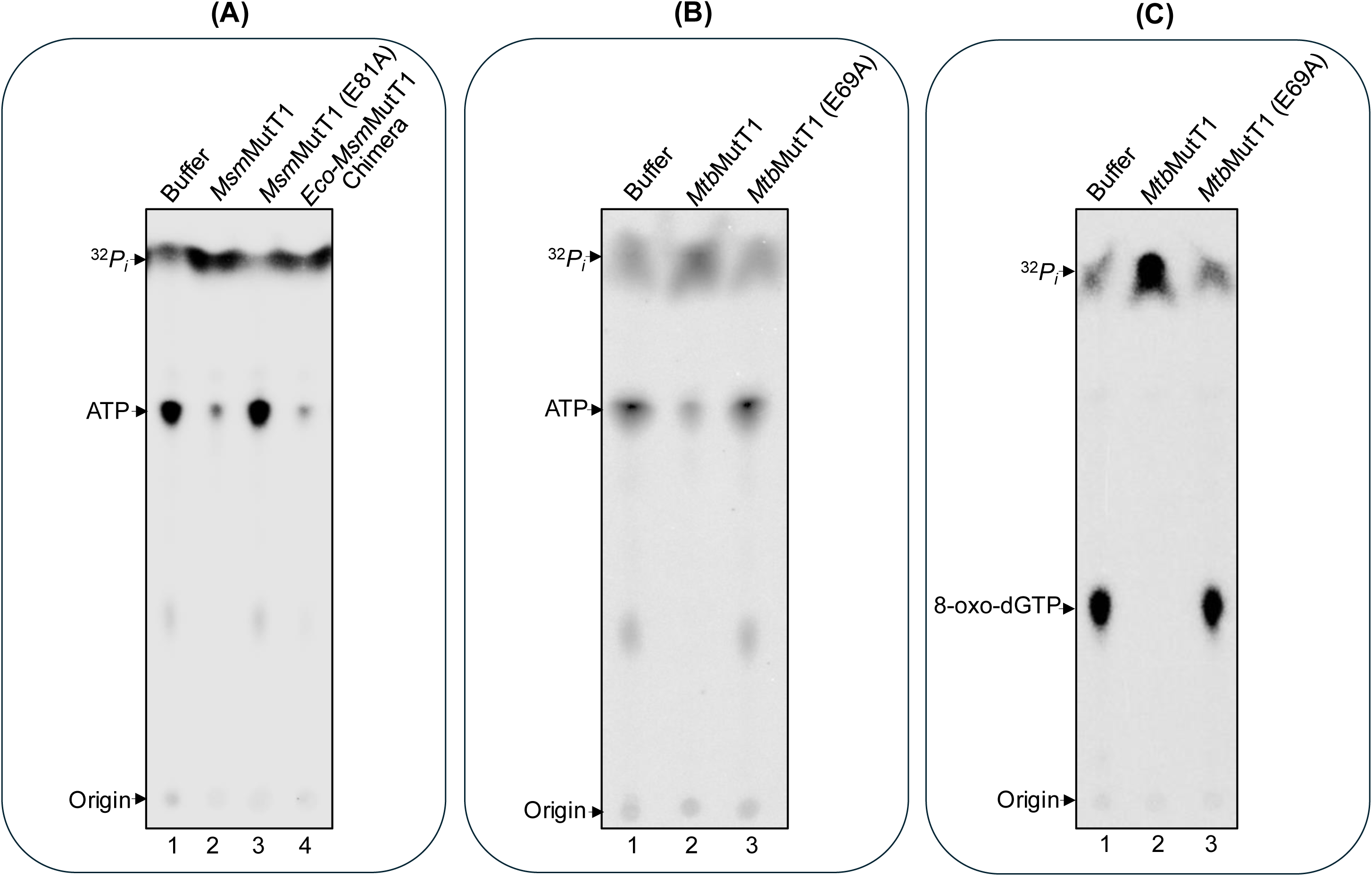
Regulation of NDK activity in vitro. **(A) ADP to ATP conversion by *Msm*NDK.** *Msm*NDK-*Pi* (1 μg) was incubated with buffer (lane 1), 2 μg *Msm*MutT1 (lane 2), 2 μg *Msm*MutT1 E81A (lane3), or 2 μg *Eco-Msm*MutT1 chimera (lane 4) at 30 °C for 1 h followed by addition of ADP as described in Materials and Methods. For percent conversion of ADP to ATP upon incubation of ADP with *Msm*NDK + buffer, *Msm*NDK + *Msm*MutT1, *Msm*NDK + *Msm*MutT1 E81A, or *Msm*NDK + *Eco-Msm*MutT1 chimera, refer to Fig. S11A *panel* i. For the percentage of the free phosphate that resulted from incubation of *Msm*NDK with buffer, *Msm*MutT1, *Msm*MutT1 E81A, or *Eco-Msm*MutT1 chimera, refer to Fig. S11A, *panel* ii. **(B) ADP to ATP conversion by *Mtb*NDK**. The *Mtb*NDK-*Pi* (1 μg) was incubated with buffer (lane 1), 2 μg *Mtb*MutT1 (lane 2), or 2 μg *Mtb*MutT1 E69A (lane 3) at 30 °C for 1 h followed by addition of ADP as described in Materials and Methods. For the percent conversion of ADP to ATP upon incubation of ADP with *Mtb*NDK + buffer, *Mtb*NDK + *Mtb*MutT1, or *Msm*NDK + *Msm*MutT1 E69A refer to Fig. S11B, *panel* i. For the percent of the free phosphate resulting from incubation of *Mtb*NDK with buffer, *Mtb*MutT1, or *Mtb*MutT1 E69A refer to Fig. S11B, *panel* ii. **(C) 8-oxo-dGDP to 8-oxo-dGTP conversion by *Mtb*NDK**. The *Mtb*NDK-*Pi* (1 μg) was incubated with buffer (lane 1), 2 μg *Mtb*MutT1 (lane 2), or 2 μg *Mtb*MutT1 E69A (lane 3) for 1 h followed by addition of 8-oxo-dGDP as described in Materials and Methods. For percent conversion of the 8-oxo-dGDP to 8-oxo-dGTP refer to Fig. S11C, *panel* i. For the percent of the free phosphate resulting from incubation of *Mtb*NDK with buffer, *Mtb*MutT1 or *Mtb*MutT1 E69A refer to Fig. S11C, *panel* ii. Three technical replicates were used in these experiments.

### MutT1 regulates the NDK and affects its downstream activity *in vivo*

8-oxo-dGTP misincorporation in DNA causes AT to CG transversions. *Msm*MutT1 hydrolyses 8-oxo-(d)GTP to 8-oxo-(d)GDP and prevents its misincorporation in DNA. However, overexpression of NDK in *E. coli* CC101 Δ*mutT* strain results in increased frequency of A to C mutations because of its role in conversion of 8-oxo-dGDP to 8-oxo-dGTP [19]. As MutT1 can dephosphorylate NDK-*Pi* and prevent it from converting 8-oxo-dGDP to 8-oxo-dGTP, we investigated the physiological relevance of this activity by utilizing *E. coli* CC101 Δ*mutT*Δ*ndk*::*kan*. The β-galactosidase (LacZ) gene in the CC101 strain is inactive because of an amber mutation (G to T) at E461 codon (GAG to TAG) in the active site. For the strain to grow on lactose (as sole carbon source) it requires a specific reversion mutation of A to C (or T to G) at this site. To investigate the effect of MutT1 on NDK, we carried out assays using the strains harbouring pACDH*Eco*NDK.

We transformed the CC101 Δ*mutT*Δ*ndk*::*kan*/pACDH*Eco*NDK strain with the plasmid borne copies of *Eco*MutT, *Msm*MutT1, and *Msm*MutT1 (E81A), and determined the reversion frequencies (Fig. 6A). As expected, CC101Δ*mutT*Δ*ndk*::*kan*/pACDH*Eco*NDK showed a higher mutation frequency of 1.8×10^-6^ compared to the controls CC101/pACDH, CC101 Δ*mutT*Δ*ndk*::*kan*/pACDH*Eco*NDK/pBAD*Eco*MutT (undetectable mutation frequency) and CC101Δ*mutT*Δ*ndk*::*kan*/pACDH with mutation frequency of 7.5×10^-7^. The mutation frequency decreased to 1×10^-6^ when *CC101*Δ*mutT*Δ*ndk*::*kan/*pACDH*EcoNDK* was complemented with *Msm*MutT1. In comparison with *Msm*MutT1, complementation of *CC101*Δ*mutT*Δ*ndk*::*kan/*pACDH*EcoNDK* with *Msm*MutT1 (E81A), showed an increase in the mutation frequency (1.7×10^-6^). These observations are consistent with the in vitro observations of the MutT1 mediated regulation of NDK in converting 8-oxo-dGDP to 8-oxo-dGTP.

**Figure. 6:**
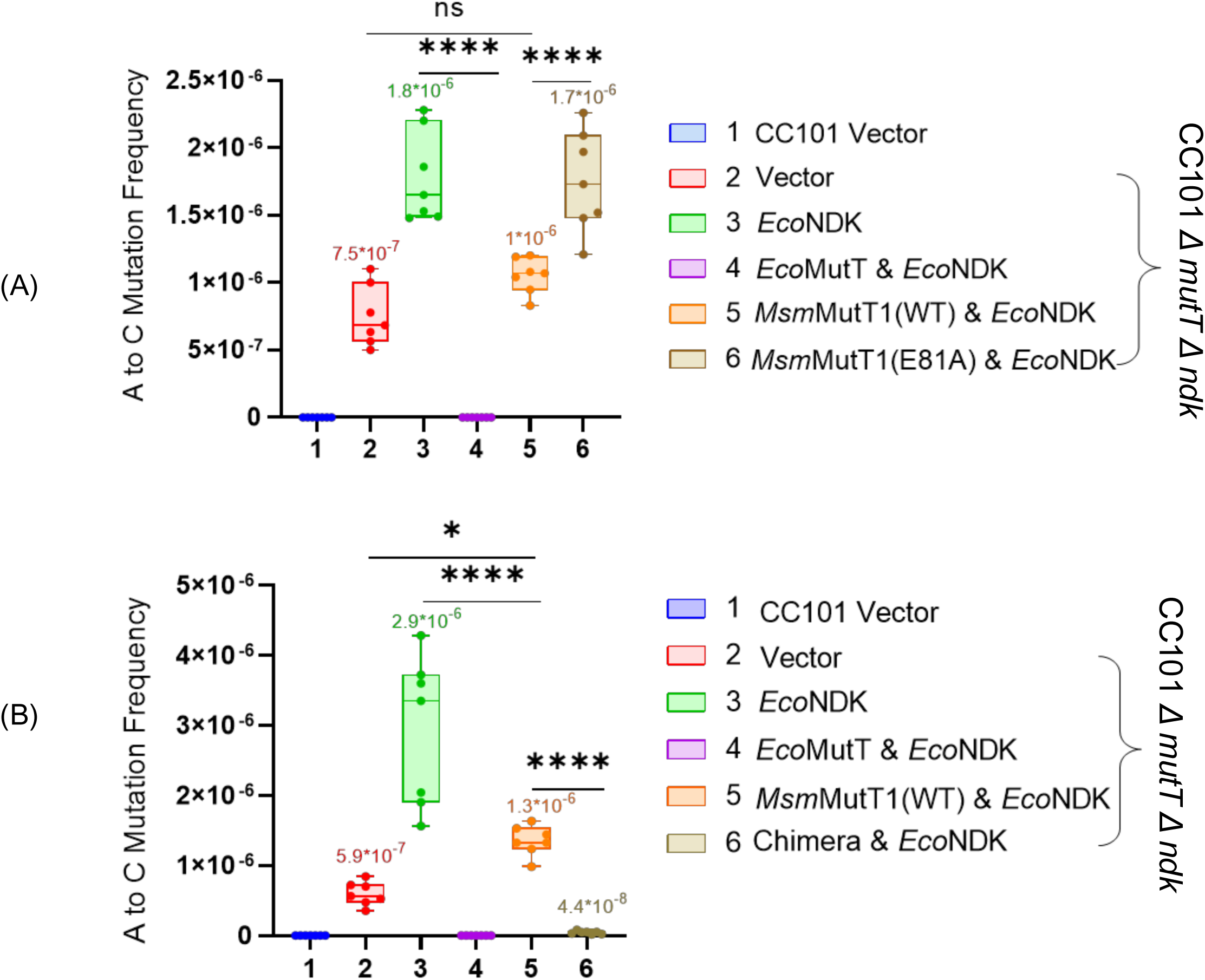
Regulation of NDK activity in vivo. Mutation frequency analysis of the Lac^+^ revertant with respect to A to C (or T to G) in *E. coli* CC101, CC101/pACDH*Eco*NDK, and its derivatives harbouring **(A)** pBADHisB (empty vector), or its recombinants harbouring *Eco*MutT, *Msm*MutT1 WT, or *Msm*MutT1 E81A. **(B)** plasmids harbouring *Eco*MutT, *Msm*MutT1 WT, or chimera. Reversion frequencies were calculated by dividing the number of colonies that appear on a minimal lactose plate by the number of colonies that appear on a minimal glucose plate. The data are represented as means ± SDs of 7 independent replicates. *p* values, ∗ *p* < 0.05; ∗∗ *p* < 0.01; ∗∗∗ *p* < 0.001 indicate significant differences between samples; ‘ns’ represent not significant. One-way ANOVA method was used to calculate *p* value.

To investigate if *Eco-Msm*MutT1 rescued CC101Δ*mutT*Δ*ndk*::*kan/*pACDH*EcoNDK* strain, we transformed the strain with plasmid borne copies of *Eco*MutT, *Msm*MutT1, and *Eco-Msm*MutT1 chimera, and determined the reversion frequencies (Fig. 6B). In this experiment too, CC101Δ*mutT*Δ*ndk*::*kan*/pACDH*Eco*NDK showed higher mutation frequency of 2.9×10^-6^ compared to the controls CC101/pACDH, CC101 Δ*mutT*Δ*ndk*::*kan*/pACDH*Eco*NDK/ pBAD*Eco*MutT (no mutation frequency was detected) and CC101Δ*mutT*Δ*ndk*::*kan/*pACDH with mutation frequency of 5.9×10^-7^. This frequency decreased to 1.3×10^-6^ when CC101Δ*mutT*Δ*ndk*::*kan/*pACDH*Eco*NDK was complemented with *Msm*MutT1. Complementation of CC101Δ*mutT*Δ*ndk*::*kan/*pACDH*Eco*NDK with *Eco-Msm*MutT1 chimera, showed an efficient decrease in the mutation frequency rate 4.4×10^-8^.

Thus, both the *in vitro* and *in vivo* observations (Figs. 5 and 6) suggest that the mycobacterial MutT1 regulates the function of NDK by dephosphorylating NDK-*Pi*.

## Discussion

8-oxo-dGTP that often results from the oxidation of dGTP, is a mutagenic nucleotide. The sources of oxidative agents can be the cellular metabolic activities or the external environments [27–30]. 8-oxo-dGTP can mis-pair with adenine and cause AT to CG transversion mutations [18, 31, 32]. Tuberculosis, caused by *M. tuberculosis*, poses an alarming threat to global public health. It results in more deaths than those caused by diseases like COVID-19 and HIV/AIDS [1]. *M. tuberculosis* encounters a high level of oxidative stress as part of the host’s innate immune response [2], and with ∼65% GC rich genome, it is highly prone to accumulate 8-oxo-dGTP in its DNA or the nucleotide pool [33]. The role of MutT proteins is important in protection against the misincorporation of the 8-oxo-dGTP, and 8-oxo-GTP into nucleic acids [19, 21, 23, 34–36]. Both *Mtb*MutT1 and *Msm*MutT1 hydrolyze 8-oxo-dGTP and 8-oxo-GTP to 8-oxo-dGDP and 8-oxo-GDP, respectively preventing their misincorporation into nucleic acids [19, 21]. Unlike *E. coli* MutT, or mycobacterial MutT2, MutT3, and MutT4, the mycobacterial MutT1 possess an extra domain (CTD) with RHG motif, histidine phosphatase [21]. Recent work from our lab revealed an unexpected role of NDK in escalating A to C mutations by converting 8-oxo-dGDP into 8-oxo-dGTP [8], raising a question if its (NDK) activity could be regulated.

In this study, comprehensive biochemical analyses reveal a protein phosphatase activity of mycobacterial MutT1 against NDK-*Pi* (Figs. 1A and 1B). This finding has been unprecedented in at least two ways. Firstly, so far, MutT proteins have been shown to possess phosphohydrolase activities against only small molecules. Mycobacterial MutT1 had also been shown to have activity on small molecules such as 8-oxo-dGTP and diadenosine polyphosphates [19, 21, 37]. This is the first study that shows any MutT to work on a protein molecule. Secondly, so far, NDK has not been shown to be regulated by dephosphorylation of its His-*Pi* residue by any proteins. However, what has been even more surprising is that based on the presence of RHG motif in MutT1 CTD, we anticipated CTD to possess this phosphatase activity. But a mutation of RHG motif to RAG did not suppress the phosphatase activity of mycobacterial MutT1 proteins against NDK-*Pi*. Instead, a mutation in the Nudix box of a critical Glu residue to Ala (E to A mutation) resulted in complete loss of the MutT1 phosphatase activity against NDK-*Pi*. It was reported that E53A mutation in Nudix hydrolase motif in *E. coli* MutT results in loss of its phosphatase activity [24]. Likewise, E162A mutation in *M. smegmatis* MutT4 Nudix hydrolase motif failed to complement the MutT4 knockout strain [23]. In *Msm*MutT1 and *Mtb*MutT1, these residues are represented by E81, and E69, respectively. Indeed, when these residues were mutated (E81A and E69A), the two MutT1 proteins lost their phosphatase activities even on *p*NPP (Figs. S2C and S3C). The His residue of the RHG motif (H170 for *Msm*MutT1 and H161 for *Mtb*MutT1) was mutated to Ala, not unexpectedly, no loss of their phosphatase activities on *p*NPP (Figs. S2C and S3C). When the activity of MutT1 mutants was tested against NDK, we recorded loss of *Msm*MutT1 E81A, and *Mtb*MutT1 E69A phosphatase activities against NDK-*Pi* (Figs. 2A and 2B), revealing the importance of Nudix hydrolase motif in dephosphorylation of even the proteins.

To confirm that only the MutT1 NTD (Nudix hydrolase motif) is responsible for its activity on NDK-*Pi*, we attempted to purify the *Msm*MutT1-NTD alone. Unfortunately, the NTD could not be purified with any reasonable purity. And, when we carried out general phosphatase activity assay using *p*NPP as substrate, we did not detect any phosphatase activity in the preparation. This could be due to the instability of the NTD sequences when expressed independent of the CTD. We then turned to *Eco*MutT and *Msm*MutT2, which possess only the Nudix hydrolase motifs domain, and they both act on 8-oxo-dGTP. Moreover, the structures of these proteins are very similar to MutT1 NTD structure (Fig. S12). When the phosphatase activities of *Eco*MutT and *Msm*MutT2 were checked against *p*NPP, high levels of phosphatase activities were detected (Fig. S5). However, when the activities of these were tested on *Eco*NDK-*Pi* and *Msm*NDK-*Pi*, respectively, neither of these dephosphorylated NDK-*Pi* (Figs. 3A and 3B). These observations then led us to ask the question of the role of the CTD of the mycobacterial MutT1 proteins. It is clear that the CTD (RHG motif) per se has no phosphatase activity on NDK-*Pi* (Figs. 2A and 2B). But the CTD may be required in some accessory ways for the NTD to act on NDK-*Pi*.

Our protein-protein interaction predictions (Fig. S6) suggested interactions between *Msm*NDK and *Msm*MutT1 CTD. To further support our hypothesis and validate the predictions, we generated a chimeric protein by replacing *Msm*MutT1 NTD with *Eco*MutT (*Eco-Msm*MutT1) (Fig. S7). We chose *Eco*MutT over *Msm*MutT2 to generate the chimeric protein because *Eco*MutT is efficient in hydrolyzing 8-oxo-dGTP into 8-oxo-dGMP [38], [39], whereas *Msm*MutT2 functions mainly as a dCTPase [34]. The chimera showed an efficient phosphatase activity on *p*NPP when compared with *Msm*MutT1 (Fig. S7). Importantly, the chimeric protein led to efficient dephosphorylation of *Eco*NDK-*Pi* or *Msm*NDK-*Pi* (Figs. 4A and 4B). These observations further support that while MutT1 NTD plays the catalytic role of dephosphorylating NDK-*Pi*, the MutT1 CTD facilitates this activity.

To understand the effect of MutT1 in regulating the downstream roles of NDK (Figs. 5A to C), we made use of the fact that NDK catalyzes reversible phosphorylation of dNDPs to dNTPs [8, 40, 41]. Our studies show that MutT1 efficiently prevents NDK mediated conversion of ADP to ATP or 8-oxo-dGDP to 8-oxo-dGTP (Figs. 5A to 5C). No effect on ADP to ATP or 8-oxo-dGDP to 8-oxo-dGTP conversion was observed when NDK-*Pi* was treated with *Msm*MutT1 E81A, or *Mtb*MutT1 E69A. The chimeric protein also behaved like the *Msm*MutT1 in preventing conversion of the ADP to ATP. Furthermore, MutT1 regulates the downstream activity of NDK not only in vitro but also in vivo (Figs. 5 and 6). Particularly, the role of the mycobacterial MutT1 proteins in downregulating the mutation frequencies in *E. coli*, suggests an important role of its evolutionarily conserved presence in mycobacteria. Based on our investigations we propose a model (Fig. 7). This model shows how MutT1 regulates the activity of NDK. In the absence of MutT1, NDK-*Pi* catalyzes the reversible conversion of 8-oxo-dGDP to 8-oxo-dGTP elevating the (A to C) mutation frequency (Fig. 7A). On the other hand, the presence of MutT1 dephosphorylates NDK-*Pi* and prevents it from converting 8-oxo-dGDP to 8-oxo-dGTP resulting in reduction of (A to C) mutation frequency (Fig. 7B).

**Figure 7:**
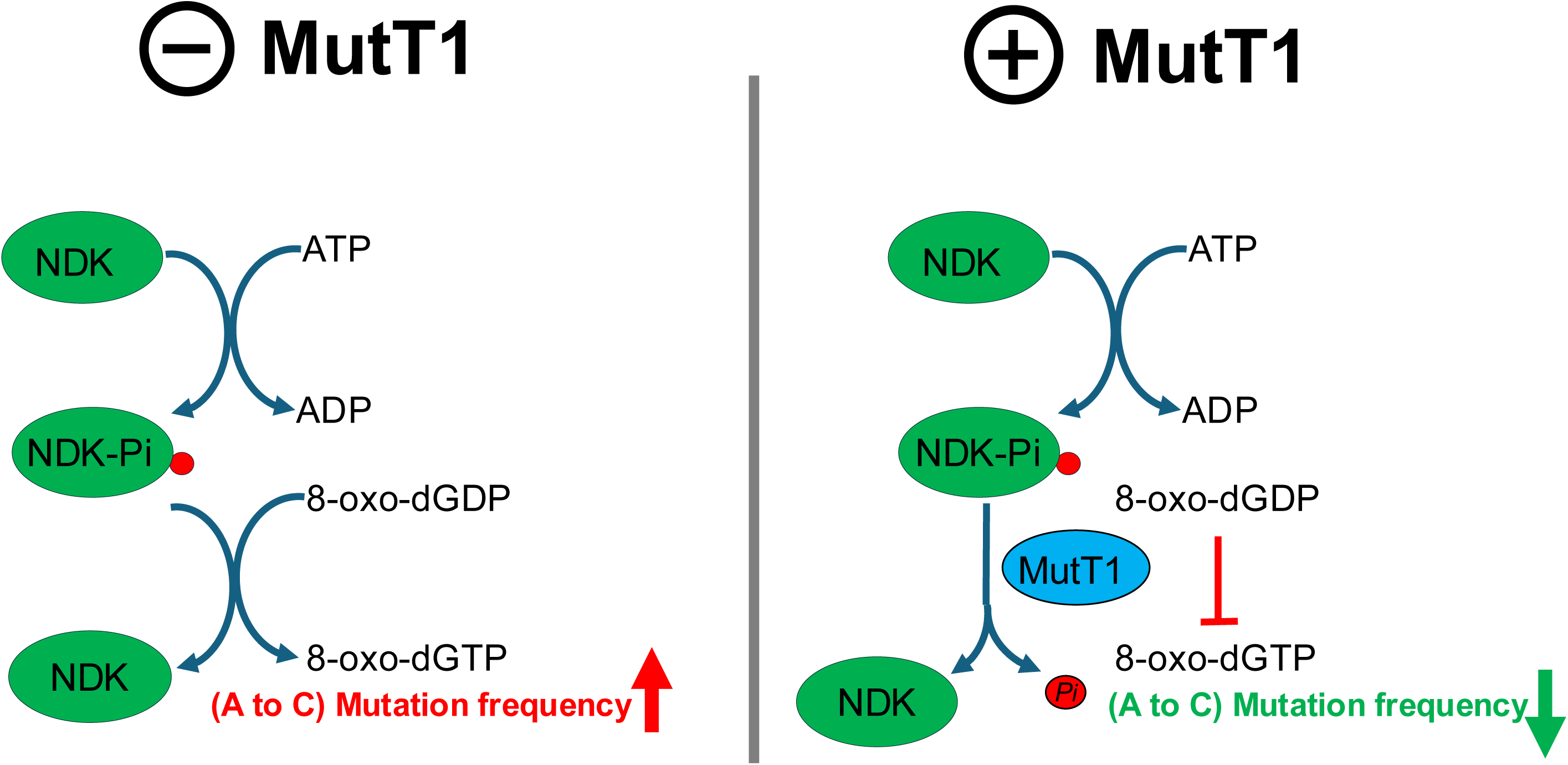
Model for the regulation of NDK activity by MutT1. **(A)** In the absence of MutT1, the active NDK-*Pi* catalyzes the conversion of 8-oxo-dGDP to 8-oxo-dGTP elevating A to C mutation frequency. **(B)** MutT1 dephosphorylates NDK-*Pi* preventing it from converting 8-oxo-dGDP to 8-oxo-dGTP resulting in reduction of A to C mutation frequency.

Finally, as we only tested the effect of MutT1 in maintaining the genome stability by regulating the activity of NDK, we cannot rule out the possibility that MutT1 might also regulate the other reported roles of NDK such as regulation of host defense mechanisms, interaction of NDK with FtsZ, bacterial growth, and signal transduction pathways [12, 42, 43]. Currently, we are in the process of understanding the enzymatic activity of (RHG motif) in the CTD of MutT1.

## Materials and methods

### Bacterial strains, plasmids, enzymes, nucleotides and DNA oligomers

Strains and plasmids used in this study are described in Table 1. DNA oligomers are listed in Table S1. Restriction enzymes (RE) and other DNA modifying enzymes used for cloning were ordered from New England Biolabs and Thermo Fisher Scientific. The 8-oxo-dGDP was purchased from Jena Bioscience, Germany. Media components were bought from BD Difco (Franklin Lakes, NJ).

**Table 1:** Description of strains and plasmids used in the study.

| Strain/Plasmid | Genotype | Reference/<br>Source |
| --- | --- | --- |
| *E. coli* TG1 | *supE Δ(lac-proAB) hsdΔ5F' [traD36 proAB+lacIq lacZΔM15]* | Novagen |
| *E. coli* BL21 Rosetta (DE3) | F' *ompT hsdSB(rB-mB-) gal dcm* (DE3) pRARE (CamR) | Novagen |
| JW0097 *ΔmutT::kan* | *Δ mutT790::kan*, LAM2 rph-1 | [46] |
| *E. coli* M15/pREP4 | *E. coli* strain with deletion of M15 in the lacZ gene. pREP4 codes for a repressor for leaky expression prevention. | [47] |
| *E. coli* CC101 | F', *ara-600, Δ(gpt-lac)5, λ-, relA1, spoT1, thiE1*, F128- (used to screen for A to C or T to G mutation) | [48] |
| *E. coli* CC101Δ*ndk::kan ΔmutT* | CC101 *ΔmutT* cured strain, where *ndk* has been disrupted by introducing *kan* cassette | [8] |
| *M. smegmatis* *mc<sup>2</sup>155* MSMEG_2390(*mutT1*)::*hyg* | *mc<sup>2</sup>155* strain where *mutT1* gene has been disrupted with *hyg* cassette | Unpublished data |
| pBADHisB (pBAD) | *E. coli* expression vector harboring ColE1 ori of replication, *amp<sup>R</sup>* marker, and arabinose inducible promoter | Invitrogen |
| pBAD *Eco* *mutT* | pBAD plasmid contains the *Eco_mutT* gene cloned (NcoI, NheI) under arabinose promoter | [34] |
| pTrcNdeHis-MsMutT1 | pTrcNdeHis plasmid with *Msm mutT1* gene cloned in its NdeI/HindIII site | [21] |
| pBAD *Msm* MutT1 | *Msm mutT1* was digested from pTrcNdeHis-MsMutT1 with (NcoI & HindIII) and subcloned into pBADHisB | This study |
| pACDH | A low copy number plasmid with ACYC ori of replication, compatible with the ColE1 origin of replication (TetR) | [49] |
| pACDH *Eco* *ndk* | pACDH plasmid contains the *Eco_ndk* gene cloned (NcoI, HindIII) under IPTG promoter | This study |
| pJAM_His | Shuttle vector harboring ColE1 (*E. coli*) and pAL5000 (mycobacteria) origin of replications, *kan<sup>R</sup>* marker, and mycobacterial acetamidase promoter | [50] |
| pJAM_ *Msm* *mutT1* | *Msm mutT1* was cloned into pJAM_His between BamHI and XbaI sites. | This study |
| pJAM_ *Msm* *mutT1* E81A | pJAM with *Msm mutT1* E81A mutant | This study |
| pJAM_ *Msm* *mutT1* H170A | pJAM with *Msm mutT1* H170A mutant | This study |
| pJAM_ *Mtb* *mutT1* | *Mtb mutT1* was cloned into pJAM_His in XbaI site. | This study |
| pJAM_ *Mtb* *mutT1* E69A | pJAM with *Mtb mutT1* E69A mutant | This study |
| pJAM_ *Mtb* *mutT1* H161A | pJAM with *Mtb mutT1* H161A mutant | This study |
| pTrcMsm-*mutT2* | pTrc99c plasmid with *Msm-mutT2* cloned in its NcoI and HindIII sites (from pET14b*Msm-mutT2*) | [34] |
| pPROEX-HTb *Eco-MsmMutT1* chimera | Chimera (*Eco mutT* + *Msm mutT1* CTD) was cloned into pProEx-Htb between BamHI and XhoI sites. | This study |
| pBAD_ *Eco_ndk* | A pBAD plasmid having *Eco_ndk* cloned (NcoI, BglII) under arabinose promoter | [8] |
| pPROEX-HTb *Msm* *ndk* | *Msm ndk* gene was cloned into pProEx-Htb between NcoI and HindIII sites. | This study |
| pQE30*Mtb-ndk* | *Mtb ndk* gene was cloned into pQE30 between BamHI and HindIII | [51] |
| pBAD *Msm* *mutT1* E81A | pBAD plasmid with *Msm mutT1* E81A mutant | This study |
| pBADEco-*MsmMutT1* chimera | pPROEX chimera was digested with NcoI & XhoI and chimera was subcloned into pBADHisB. | This study |

### Media, and growth conditions

*E. coli* strains were grown in Luria-Bertani (LB) medium. For growth on solid surface, 1.6% agar was added to LB medium. Minimal medium plates used for (A to C) reversion assays contained 1× M9 salts (47 mM Na_2_HPO_4_, 22 mM KH_2_PO_4_, 8.5 mM NaCl, and 18.6 mM NH_4_Cl), 2 mM MgSO_4_, 0.1 mM CaCl_2_, 5 μg/ml thiamine, 0.2% glucose (or lactose), and 1.6% agar. Media were supplemented with ampicillin (Amp), hygromycin (Hyg), kanamycin (Kan), and tetracycline (Tet) as required at concentrations of 100 μg/ml, 150 μg/ml, 25 μg/ml, and 7.5 μg/ml, respectively, unless mentioned otherwise. For culturing *M. smegmatis* strains, LB media containing 0.2% tween 80 (v/v) (LBT) was used. For growing *M. smegmatis* strains on solid media, LB containing 0.04% tween 80 (v/v) was supplemented with 1.5% agar (w/v). Media were supplemented with Kan (50 μg/ml) and Hyg (50 μg/ml), as required.

### Cloning of *Msm*MutT1

*Msm*MutT1 (MSMEG_2390) open reading frame (ORF) was amplified from the *M. smegmatis* mc^2^155 genomic DNA using *Msm* mutT1 BamH1 forward primer (Fp) and *Msm* mutT1 Xba1 reverse primer (Rp) primers. The PCR was incubated at 94 °C for 5 min followed by 30 cycles of incubations at 94 °C for 1 min, 63 °C for 30 s, and 72 °C for 2 min, and then 72 °C for 5 min. The PCR product was purified, digested with BamHI and XbaI, and cloned into similarly digested pJAM_His. The clones were confirmed by RE digestions and DNA sequencing.

### Cloning of *Mtb*MutT1

*Mtb*MutT1 (Rv2985) ORF was amplified from *M. tuberculosis* H37Rv genomic DNA using *Mtb* mutT1 Xba1 Fp and *Mtb* mutT1 Xba1 Rp. The PCR was incubated at 94 °C for 5 min followed by 30 cycles of heatings at 94 °C for 1 min, 60 °C for 30 s, and 72 °C for 2 min, and then at 72 °C for 5 min. The PCR product was purified, digested with XbaI, cloned into similarly digested pJAM_His, and confirmed by RE digestions and DNA sequencing.

### Cloning of *Msm*NDK

The ORF of *Msm*NDK (MSMEG_4627) was amplified from *M. smegmatis* mc^2^155 genomic DNA using *Msm* ndK-Ncoi-Fp and *Msm* ndK-Hindiii-Rp. The PCR was incubated at 94 °C for 5 min followed by 30 cycles of incubations at 94 °C for 1 min, 58 °C for 30 s, and 72 °C for 1 min, and then at 72 °C for 5 min. The purified PCR product was digested with NcoI and HindIII and cloned into similarly digested pProEx-Htb. The clone was confirmed by RE digestions and DNA sequencing.

### Generation of MutT1 mutants

*Msm*MutT1 (E81A or H170A), and *Mtb*MutT1 (E69A or H161A) were generated by site-directed mutagenesis (SDM), using pJAM*Msm*MutT1 and pJAM*Mtb*MutT1 templates, respectively. The primer combinations of *Msm* mutT1 E81A Fp and *Msm* mutT1 E81A Rp; *Msm* mutT1 H170A Fp and *Msm* mutT1 H170A Rp; *Mtb* mutT1 E69A Fp and *Mtb* mutT1 E69A Rp; *Mtb* mutT1 H161A Fp, and *Mtb* mutT1 H161A Rp were used. The reactions with 100 ng template DNA, 10 pmols each of the forward and reverse primers, 250 μM dNTPs, 1× Q5 polymerase buffer, and Q5 polymerase enzyme (NEB #M0491) were heated at 98 °C for 1 min followed by 20 cycles of incubations at 98 °C for 20 s, 68 °C for 30 s, and 72 °C for 6 min and final extension at 72 °C for 10 min. The PCR product was digested with DpnI (Thermo Scientific™ # ER1702) and used to transform *E. coli* TG1. Plasmids from the transformants were confirmed for SDM by DNA sequencing.

### Chimeric *Eco-Msm*MutT1 generation

The chimera generation was done in four steps. Firstly, *Eco*MutT was amplified from pBAD *Eco*MutT using the *Eco* mutT BamHI Fp and *Eco* mutT chimera Rp. The PCR was incubated at 94 °C for 3 min followed by 30 cycles of incubations at 94 °C 1 min, 65 °C 30 s, 70 °C 1 min 30 s followed by a final extension at 70 °C for 5 min. Secondly, *Msm*MutT1 CTD was amplified from pJAM *Msm*MutT1 using *Msm* mutT1 CTD Fp and *Msm* mutT1 CTD XhoI Rp. The PCR was incubated at 94 °C for 3 min followed by 30 cycles of incubations at 94°C for 1 min, 65 °C for 30 s, 70°C for 1 min 30 s followed by a final extension at 70 °C for 5 min. Thirdly, for the overlapping PCR, PCR products *Eco*MutT and *MsmMutT1*-CTD were used to generate a full-length fragment. The PCR reaction was incubated at 94 °C for 3 min followed by 12 cycles of incubations at 94 °C 1 min, 56 °C 20 min, 70 °C 1 min followed by a final extension at 70 °C 10 min. Fourthly, for the chimera cloning, the PCR product from the third step was used as template to amplify the chimeric gene using *Eco* mutT BamHI Fp and *Msm* mutT1 CTD XhoI Rp. The PCR was incubated at 98 °C for 30 s followed by 30 cycles of incubations at 98 °C for 10 s, 65 °C for 30 s, 72 °C for 35 s and a final extension at 72 °C for 2 min. The PCR product was purified, digested with BamHI and XhoI and cloned into similarly digested pProEx-Htb. The clones were confirmed by RE digestions and DNA sequencing.

### Purification of MutT1 proteins

MutT1 proteins were purified from *M. smegmatis* mc^2^155 *ΔmutT1* strain. Briefly, the expression plasmids containing MutT1 constructs, were electroporated into mc^2^155 *ΔmutT1*. An isolated colony was inoculated in LB with 0.2% tween 80 and Kan. Inoculum (1% of the saturated culture) was sub-cultured into 3 L LB with 0.2% tween 80 and Kan. At OD_600_ of 0.6, protein expression was induced with 1% acetamide and the culture was further incubated at 30 °C for 9 h. The cells were pelleted by centrifugation, resuspended in buffer A (20 mM Tris-HCl, pH 8.0, 1 M NaCl, 10% (v/v) glycerol, 20 mM imidazole and 2 mM β-mercaptoethanol), lysed by sonication and ultracentrifuged at 26K rpm at 4 °C for 2 h (using optima XPNs Ultra Centrifuge, Beckman Coulter). The clarified lysate was loaded onto pre-equilibrated Ni-NTA column, and washed with wash buffer (1 M NaCl, 20 mM Tris-HCl pH 8, 10% (v/v) glycerol, 2 mM β-mercaptoethanol and 40 mM imidazole). The proteins were eluted with a gradient of 40 to 1000 mM imidazole in wash buffer lacking imidazole. The fractions containing the desired protein were pooled, concentrated, and loaded onto Superdex-75 column, and eluted with gel filtration buffer (1 M NaCl, 20 mM Tris-HCl pH 8, 10% (v/v) glycerol and 2 mM β-mercaptoethanol). The fractions containing the purified proteins were pooled and dialyzed against dialysis buffer (750 mM NaCl, 20 mM Tris-HCl pH 8, 10% (v/v) glycerol, and 2 mM β-mercaptoethanol), followed by dialysis in storage buffer (500 mM NaCl, 20 mM Tris-HCl pH 8, 50% glycerol, and 2 mM β-mercaptoethanol) and stored at -20 °C.

### Purification of *Msm*NDK

*Msm*NDK was expressed and purified using pPROEX-HTb *Msm*NDK. Briefly, *E. coli* M15 was transformed with pPROEX-HTb *Msm*NDK. A single isolated colony was used to inoculate for starter culture in 25 ml LB containing Kan and Amp. Saturated culture (1%) was used to inoculate 2 L LB with Kan and Amp, and incubated under shaking at 37 °C. When the culture reached an OD_600_ of ∼ 0.6, cells were induced with 1 mM IPTG and incubated further at 37 °C for 4 h. Cells were pelleted at 8K RPM at 4 °C for 5 min using Kubota 6500 centrifuge and resuspended in buffer A (500 mM NaCl, 50 mM Tris-HCl pH 8, 10% v/v glycerol, 10 mM imidazole, 2 mM β-mercaptoethanol and 1 mM PMSF). Resuspended cells were sonicated (2 s on / 2 s off cycles for 1 min at 35% amplitude), centrifuged at 13K at 4°C for 30 min. The supernatant was then subjected to ultracentrifugation at 26K RPM at 4 °C for 2 h (using optima XPNs Ultra Centrifuge, Beckman Coulter). Cell-free extract so prepared was loaded onto 1 ml pre-equilibrated Ni-NTA column. The column was washed with 50 mM imidazole. Elution of proteins was done using a gradient of 10 mM to 1 M imidazole in 30 ml buffer A (without imidazole). Fractions were loaded on 12% SDS-PAGE. The fractions containing the desired protein were pooled, concentrated, and loaded onto Superdex-75 column, and eluted with gel filtration buffer (500 mM NaCl, 20 mM Tris-HCl pH 8, 10% (v/v) glycerol and 2 mM β-mercaptoethanol). Fractions enriched with the protein were dialysed against 50% glycerol, 150 mM NaCl, 50 mM Tris-HCl pH 8, and stored in -20 °C.

### Purification of *Eco-Msm*MutT1 chimera

*E. coli* BL21 was transformed with pPROEX-HTb-*Eco-Msm*MutT1, and an isolated colony was inoculated in 25 mL LB containing Amp. Saturated culture (1%) was used to inoculate 2 L LB, and incubated under shaking at 37 °C for 3 h. Cells were induced with 1 mM IPTG and grown further at 25 °C for 8 h. Cells were pelleted at 8K RPM, at 4 °C for 5 min, resuspended in buffer A (1000 mM NaCl, 50 mM Tris-HCl pH 8, 10% (v/v) glycerol, 10 mM imidazole, 2 mM β-mercaptoethanol and 1 mM PMSF), sonicated (2 s on / 2 s off cycles, for 1 min at 35% amplitude), and centrifugated at 13K, 4 °C for 30 min. The supernatant was subjected to ultracentrifugation at 26K rpm at 4 °C for 2 h (using optima XPNs Ultra Centrifuge, Beckman Coulter). Cell lysate was loaded onto 1 ml pre-equilibrated Ni-NTA column. After loading, the column was washed with 50 mM imidazole. Proteins were eluted using a gradient of 10 mM to 1 M imidazole in 30 ml buffer A without imidazole. Fractions were analysed on 12% SDS-PAGE, and those containing the desired protein were pooled, concentrated, loaded onto Superdex-75 column, and eluted with 1 M NaCl, 20 mM Tris-HCl pH 8, 10% (v/v) glycerol and 2 mM β-mercaptoethanol. Fractions enriched for the desired protein were dialysed against 50% glycerol, 500 mM NaCl, 50 mM Tris-HCl, pH 8 and stored in -20 °C.

### Purification of *Mtb*NDK, *Eco*NDK, *Eco*MutT, and *Msm*MutT2

The *Mtb*NDK, *Eco*NDK, *Eco*MutT, and *Msm*MutT2 were purified by Ni–NTA affinity chromatography followed by size-exclusion chromatography using a Superdex 75 column. The culture growth and protein purifications were done essentially as described [8], [34], [44].

### General phosphatase activity assay

The assays were performed as described [45]. Briefly, purified protein (1 μg) was incubated with 10 mM *para*-Nitrophenylphosphate (*p*NPP) in a 50 μl reactions consisting of 50 mM Tris-HCl pH 7.5, 25 mM NaCl, and 100 mM MgCl_2_ at 37 °C for 30 min. The reactions were stopped by adding 150 μl 1N NaOH. The absorbance at 405 nm was read in Tecan plate reader.

### Preparation of NDK-*Pi* and its use in dephosphorylation assays

Briefly, 1 μg of purified NDK was incubated in kinase buffer (50 mM Tris-HCl, pH 8.0, 50 mM KCl, 10 mM MgCl_2_) containing 1 μCi of γ-^32^P-ATP (3000 Ci/mM) at 30 °C for 1 h to generate NDK-*Pi*. Further incubations with 1 to 2 μg MutT1 proteins were done for 1 h at 30 °C. The reactions were terminated by adding 1× SDS-PAGE sample buffer and analysed on 12% SDS-PAGE (because the phospho-His is labile to heat, the samples were not heated in sample buffer). After electrophoresis, the gel was exposed to phosphor imager screen (Fujifilm Bas cassette2, Japan) for 4 h followed by imaging with Typhoon 9210 phosphor imager (GE Healthcare, USA) and Azure Sapphire (Azure Biosystems, USA).

### ADP to ATP, or 8-oxo-dGDP to 8-oxo-dGTP conversion assay

Briefly, purified NDK (1 μg) was incubated in kinase buffer (50 mM Tris-HCl, pH 8.0, 50 mM KCl, 10 mM MgCl_2_) containing 1 μCi of γ-^32^P-ATP (3000 Ci/mM) at 30 °C for 1 h. Further incubation were with 1 to 2 μg MutT1 proteins for 1 h at 30 °C. ADP, or 8-oxo-dGDP were added to the reaction to 1 mM and incubated at 30 °C for 5 min. The reaction was stopped by 2% (final concentration) HCOOH. The reaction aliquots (2.5 μl) were spotted on PEI Cellulose F plate (MERCK # 105579), developed with 1.5 M KH_2_PO_4_ (pH 3.4), dried, exposed to phosphor imager screen (Fujifilm Bas cassette2, Japan) for 4 h, and imaged on Typhoon 9210 phosphor imager (GE Healthcare, USA) and Azure Sapphire (Azure Biosystems, USA).

### A to C reversion assay

*E. coli* CC101 and CC101Δ*mutTΔndk* were transformed with relevant plasmids, and grown on LB agar containing 50 μg/ml 5-bromo-4-chloro-3-indolyl β-D-galactopyranoside (X-Gal) and suitable antibiotics. Isolated white colonies were inoculated into 2 ml LB having 0.005% arabinose, 100 μM IPTG, desired antibiotics and grown at 37 °C for 16 h. OD_600_ was normalized to 1.5 for each replicate, and a 20 μl aliquot was spread on M9 plate containing 0.2% lactose to determine Lac^+^ revertants. Aliquots (20 μl) of 10^-5^ dilution were spread on M9 agar containing 0.2% glucose to determine the total viable counts. The plates were incubated at 37 °C for 24 h (glucose plates) and 48 h (lactose plates). The reversion frequencies were obtained by dividing the colony numbers observed on the lactose plate by those on the glucose plates for the number of replicates indicated in the figure legends.

## Data Availability Statement

The data underlying this article will be shared on reasonable request to the corresponding author.

## Acknowledgements

The authors acknowledge laboratory colleagues for their critical comments on the manuscript.

## CRediT authorship contributions statement

EAFE: Conceptualization, formal analysis, investigation, methodology, visualization, writing – original draft, writing – review & editing. UV: Conceptualization, formal analysis, methodology, supervision, funding acquisition, project administration, writing – review & editing. KR: formal analysis, investigation, methodology, writing – review & editing.

## FUNDING

This work was supported by grants from Department of Biotechnology (BT/PR50905/MED/29/1655/2023). The authors acknowledge the DBT-IISc program (BT/PR27952), and the DST-FIST level II infrastructure support. The funders had no role in study design, data collection and analysis, and decision to publish.

## CONFLICTS OF INTEREST

The authors declare no conflict of interest.

## Notes

### Competing Interest Statement

The authors have declared no competing interest.

