## Supplementary Material for "An unusual activity of mycobacterial MutT1 Nudix hydrolase domain as a protein phosphatase regulates nucleoside diphosphate kinase (NDK) function"

**Table S1. List of DNA oligomers used in the study.**

| DNA oligomer | Sequence (5' to 3') |
|---|---|
| Msm mutT1 BamH1 Fp | ACATGGGATCCGTGATGCCGGTGGAC |
| Msm mutT1 Xba1 Rp | AGGCTCTAGACTTCTCGTCGGGAGGG |
| Mtb mutT1 Xba1 Fp | ACCTTCTAGAGTGTCGATCCAGAAC |
| Mtb mutT1 Xba1 Rp | AGGCTCTAGAGGCCCGCACGTTGGC |
| Msm mutT1 E81A Fp | GCACGCGCGATCCACGAGGAGAC |
| Msm mutT1 E81A Rp | GTGGATCGCGCGTGCCGCGGCCAC |
| Mtb mutT1 E69A Fp | GTGCGGGCGATACTCGAGGAGAC |
| Mtb mutT1 E69A Rp | GAGTATCGCCCGCACCGCCCCCAC |
| Msm mutT1 H170A Fp | TGCGGGCCGGCACGGCCGGGCG |
| Msm mutT1 H170A Rp | TGCCGGCCCGCACGACGAGTACC |
| Mtb mutT1 H161A Fp | TGCGGGCTGGCACCGCGGGCAG |
| Mtb mutT1 H161A Rp | TGCCAGCCCGCACCACCAGCACC |
| Eco mutT BamHI Fp | GTCGGATCCATGAAAAAGCTGCAA |
| Eco mutT chimera Rp | CCGGTCGTTTCAGACGTTTA |
| Msm mutT1 CTD Fp | TAAACGTCTGAAACGACCGG |
| Msm mutT1 CTD Xhoi Rp | AGCGCTCGAGTTACTTCTCGTCG |
| Msm ndK-Ncoi-Fp | GCGCCATGGTGACTGAGCGGACCCTCGTA |
| Msm ndK-Hindiii-Rp | AGTAAGCTTTCAGGCGGTGGCCTCGCCGG |

Figure S1

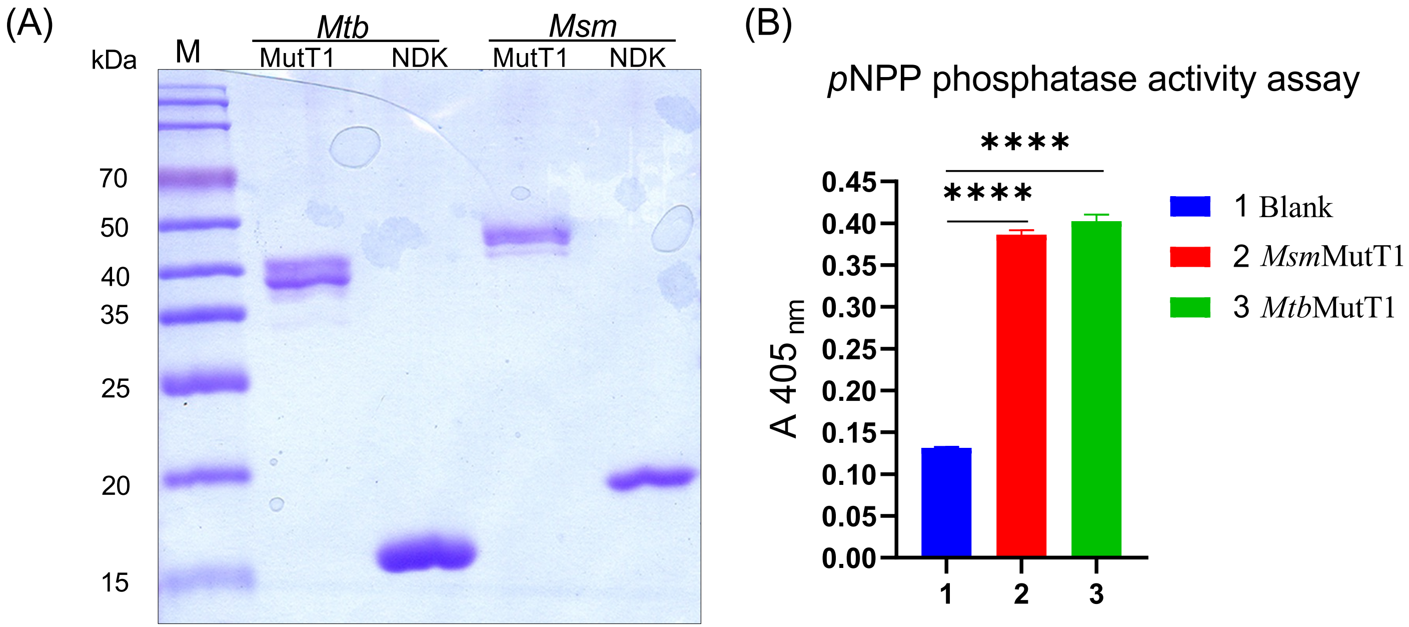


**Figure. S1: Purification of NDK and MutT1 proteins, and the general phosphatase activities of MutT1 proteins using *p*NPP**. (i) 15% SDS-PAGE gel analysis showing the quality of the purified proteins (*Mtb*MutT1, *Mtb*NDK, *Msm*MutT1 and *Msm*NDK). The calculated molecular masses of *Mtb*MutT1, *Mtb*NDK, *Msm*MutT1 and *Msm*NDK with $His_6$ tag are 37.5, 16.7, 39 and 18.1 kDa, respectively. (ii) The graph illustrates the general phosphatase activities of the MutT1 proteins using *p*NPP as a substrate. The substrate was incubated with either water (bar 1), 1 µg *Msm*MutT1 (bar 2), or 1 µg *Mtb*MutT1 (bar 3). Bars represent mean ± SD for n = 3. *p* values, $* p < 0.05$; $** p < 0.01$; $*** p < 0.001$ indicate significant differences between samples; 'ns' represent not significant. One-way ANOVA method was used to calculate *p* value.

Figure S2

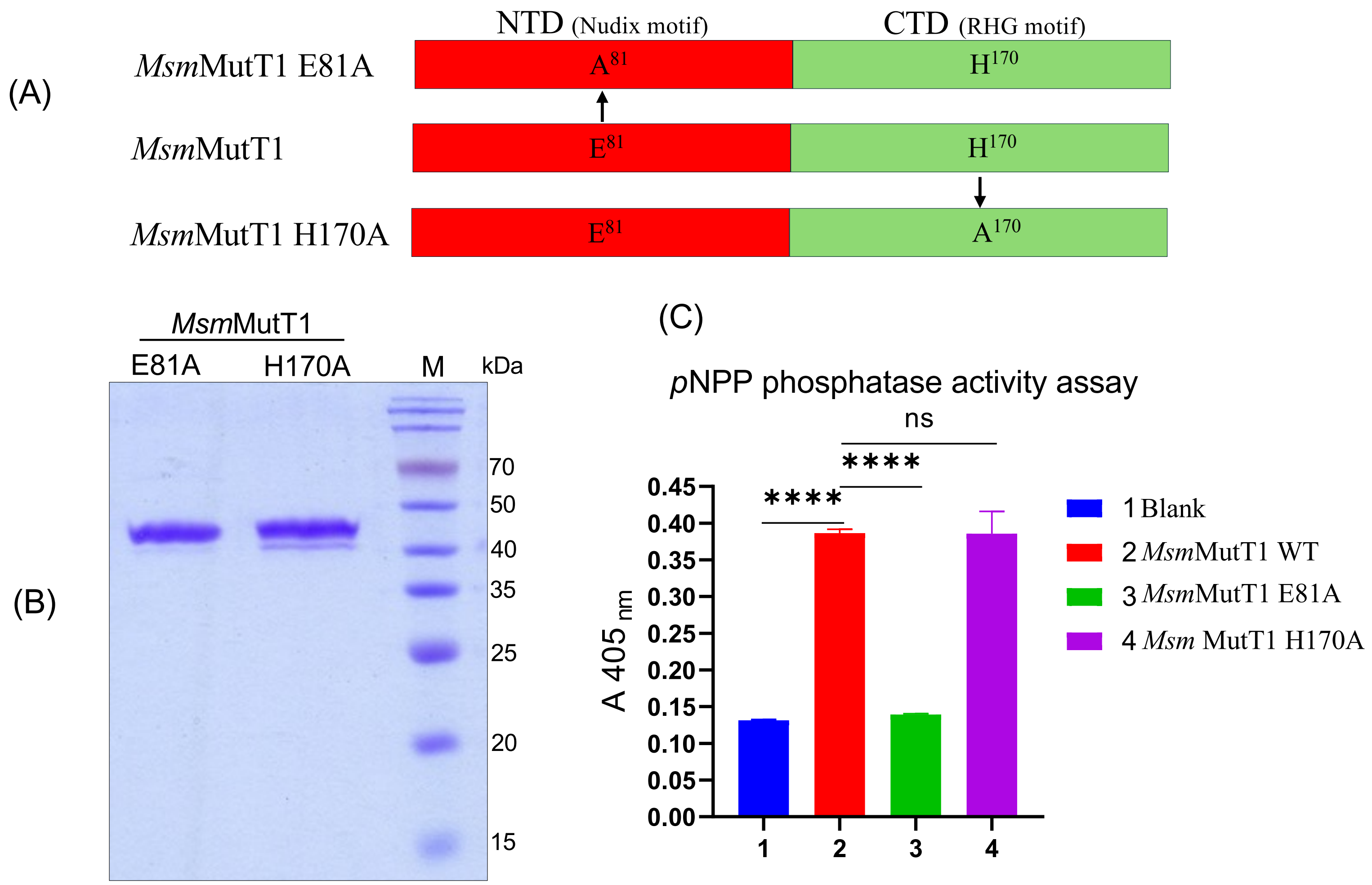


**Figure S2: Purification of *Msm*MutT1 mutant proteins and their phosphatase activity.** (A) A schematic depicting *Msm*MutT1 with its two domains, and the positions where the mutations were generated. (B) Analysis on 15% SDS-PAGE shows the quality of the purified proteins. ~3 µg of *Msm*MutT1 E81A and *Msm*MutT1 H170A were loaded into 15% SDS-PAGE gel. The calculated molecular mass of the proteins is ~39 kDa. (C) The graph illustrates the general phosphatase activity of MutT1 proteins using *p*NPP as a substrate. The substrate was incubated with either water (bar 1), 1 µg *Msm*MutT1 (bar 2), 1 µg *Msm*MutT1 E81A (bar 3) or 1 µg *Msm*MutT1 H170A (bar 4). Bars represent mean ± SD for n = 3. *p* values, $* p < 0.05$; $** p < 0.01$; $*** p < 0.001$ indicate significant differences between samples; 'ns' represent not significant. One-way ANOVA method was used to calculate *p* value.

Figure S3

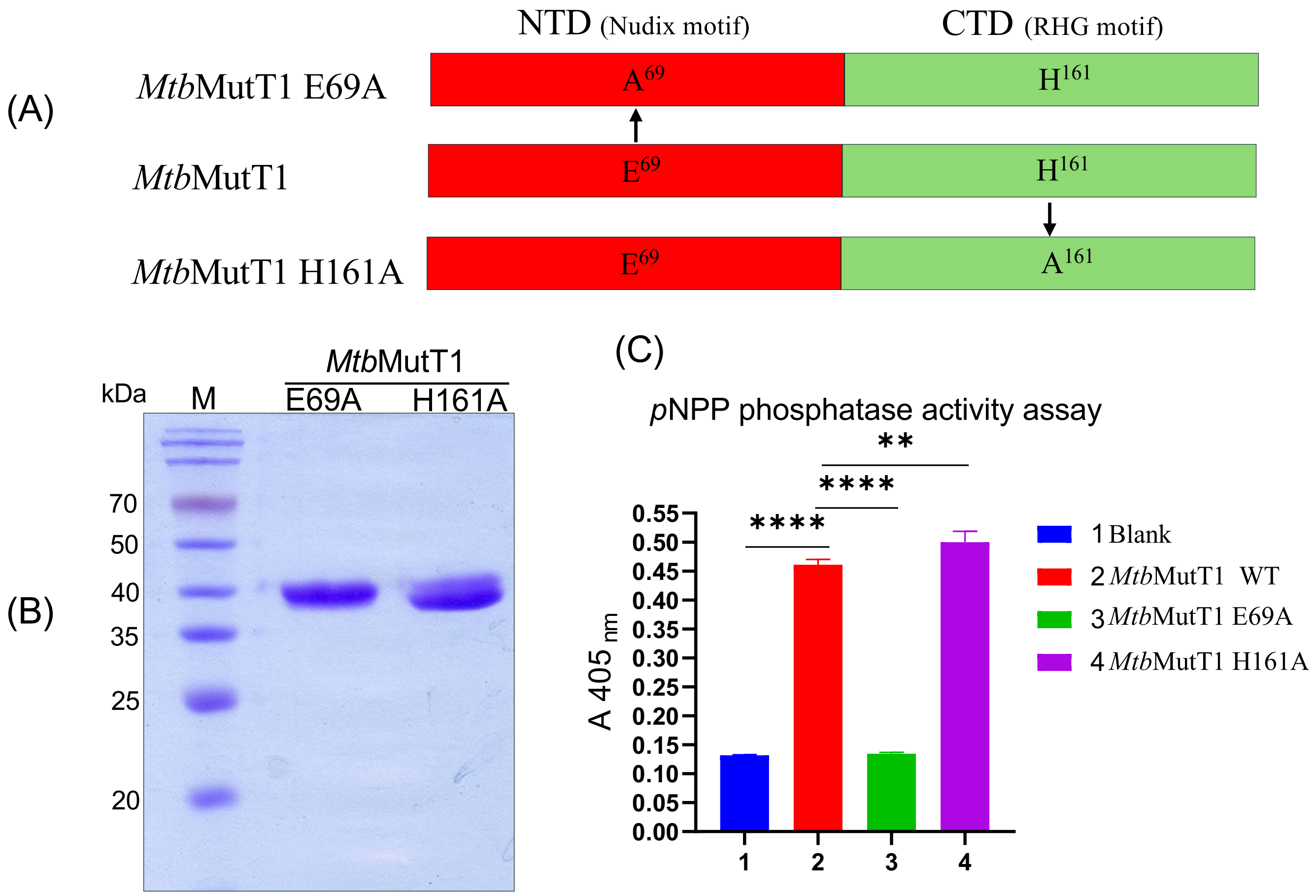


**Figure S3: Purification of *Mtb*MutT1 mutant proteins and their phosphatase activity.** (A) A schematic depicting *Mtb*MutT1 with its two domains, and the positions where the mutations were generated. (B) Analysis of the proteins on 15% SDS-PAGE gel showing the quality of the purified proteins. ~3 µg each of *Mtb*MutT1 E69A and *Mtb*MutT1 H161A were loaded into 15% SDS PAGE gel. The calculated molecular mass of the proteins is ~37 kDa. (C) The graph illustrates the general phosphatase activity of MutT1 proteins using *p*NPP as a substrate. The substrate was incubated with either water (bar 1), 1 µg *Mtb*MutT1 (bar 2), 1 µg *Mtb*MutT1 E69A (bar 3) or 1 µg *Mtb*MutT1 H161A (bar 4). Bars represent mean ± SD for n = 3. *p* values, $*$ $p < 0.05$; $**$ $p < 0.01$; $***$ $p < 0.001$ indicate significant differences between samples; 'ns' represent not significant. One-way ANOVA method was used to calculate *p* value.

Figure S4

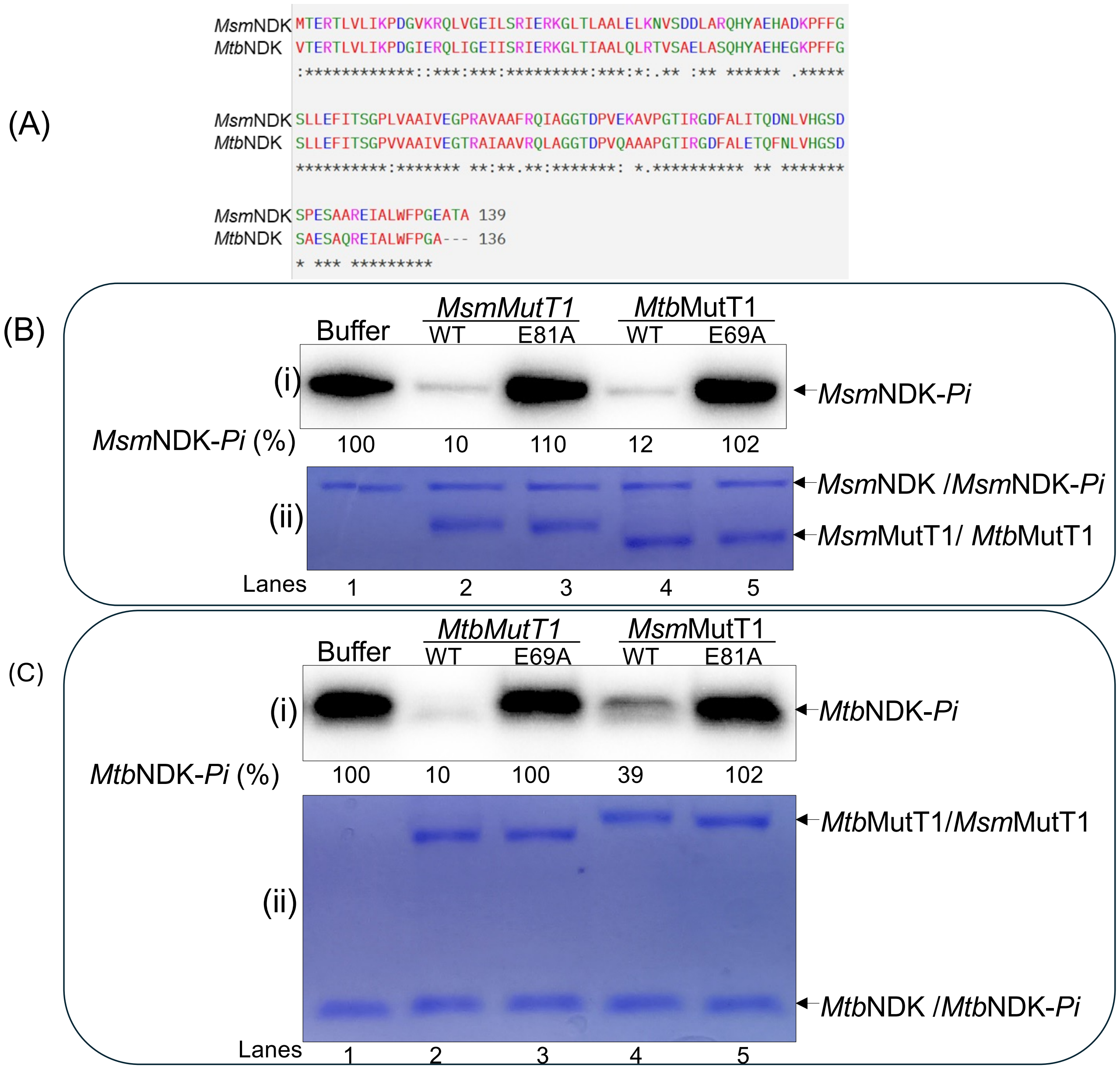


**Figure S4:Dephosphorylation of NDK-Pi with mycobacterial MutT1** (A) Sequence alignment of NDK proteins from *M. smegmatis* (*Msm*NDK) and *M. tuberculosis (Mtb*NDK*)*. Sequences were obtained from Mycobrowser website [46] and analysed using Clustal Omega Multiple Sequence Alignment (MSA) [47]. (B) Dephosphorylation of *Msm*NDK-Pi by *Msm*MutT1 (WT or E81A) and *Mtb*MutT1 (WT or E69A). *Msm*NDK-Pi (1 µg) was incubated with either buffer alone; 1 µg of *Msm*MutT1, 1 µg *Msm*MutT1 E81A, 1 µg *Mtb*MutT1 WT, or 1 µg *Mtb*MutT1 E69A (lanes 1-5, respectively). The reactions were incubated at 30 ° C for 1 h, mixed with 5 µL SDS-PAGE sample buffer and loaded (without heating) onto 12% SDS-PAGE. Gels were subjected to phosphor imaging, fixed, and stained with Coomassie brilliant blue (CBB). Panels (i) and (ii) represent autoradiogram, and CBB stained gel, respectively. Values of *Msm*NDK-Pi (%) with reference to lane 1 are shown below panel i. (C) Dephosphorylation of *Mtb*NDK-Pi by *Mtb*MutT1 (WT or E69A) and *Msm*MutT1 (WT or E81A). *Mtb*NDK-Pi (1 µg) was incubated with either buffer alone (lane 1); 1µg of *Mtb*MutT1, 1 µg *Mtb*MutT1 E69A, 1 µg *Msm*MutT1, or 1 µg *Msm*MutT1 E81A (lanes 1-5, respectively). The reactions were processed as in (A). Panels: (i) autoradiograms (ii) CBB stained gel. Values of *Msm*NDK-Pi (%) with reference to lane 1 are shown below panel i.. Because of no heating of the samples in the sample buffer, *Msm*NDK migrates slower than the expected monomeric size (also refer to Fig. S8). 2-*Msm*MutT1E81A migrates slightly faster than *Msm*MutT1 WT, while *Mtb*MutT1 migrates slower than *Mtb*MutT1 WT. This was only observed when samples are not heated.

Figure S5

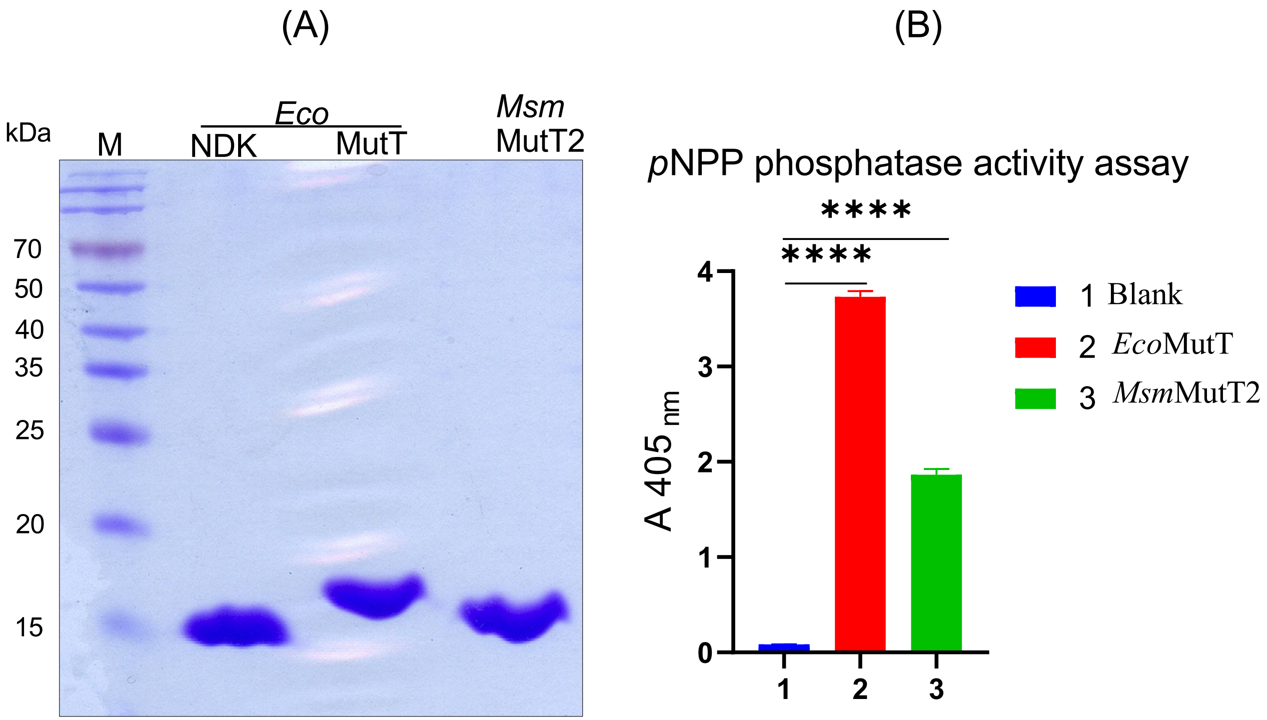


**Figure S5: Purification of *Eco*NDK, *Eco*MutT and *Msm*MutT2 proteins and the phosphatase activity of MutT proteins.** (i) Analysis on 15% SDS-PAGE gel showing quality of the purified proteins. ~5 µg of *Eco*NDK, *Eco*MutT and *Msm*MutT2, were loaded into 15% SDS PAGE gel. The calculated molecular masses of *Eco*NDK, *Eco*MutT and *Msm*MutT2 with $His_6$ tag are ~16.5, ~17 and ~14.8 kDa, respectively. (ii) The graph illustrates the general phosphatase activity of MutT proteins using *p*NPP as substrate. The substrate was incubated with either water (bar 1), 1 µg *Eco*MutT (bar 2) or 1 µg *Msm*MutT2 (bar 3). Bars represent mean ± SD for n = 3. *p* values, $* p < 0.05$; $** p < 0.01$; $*** p < 0.001$ indicate significant differences between samples; 'ns' represent not significant. One-way ANOVA method was used to calculate *p* value.

Figure S6

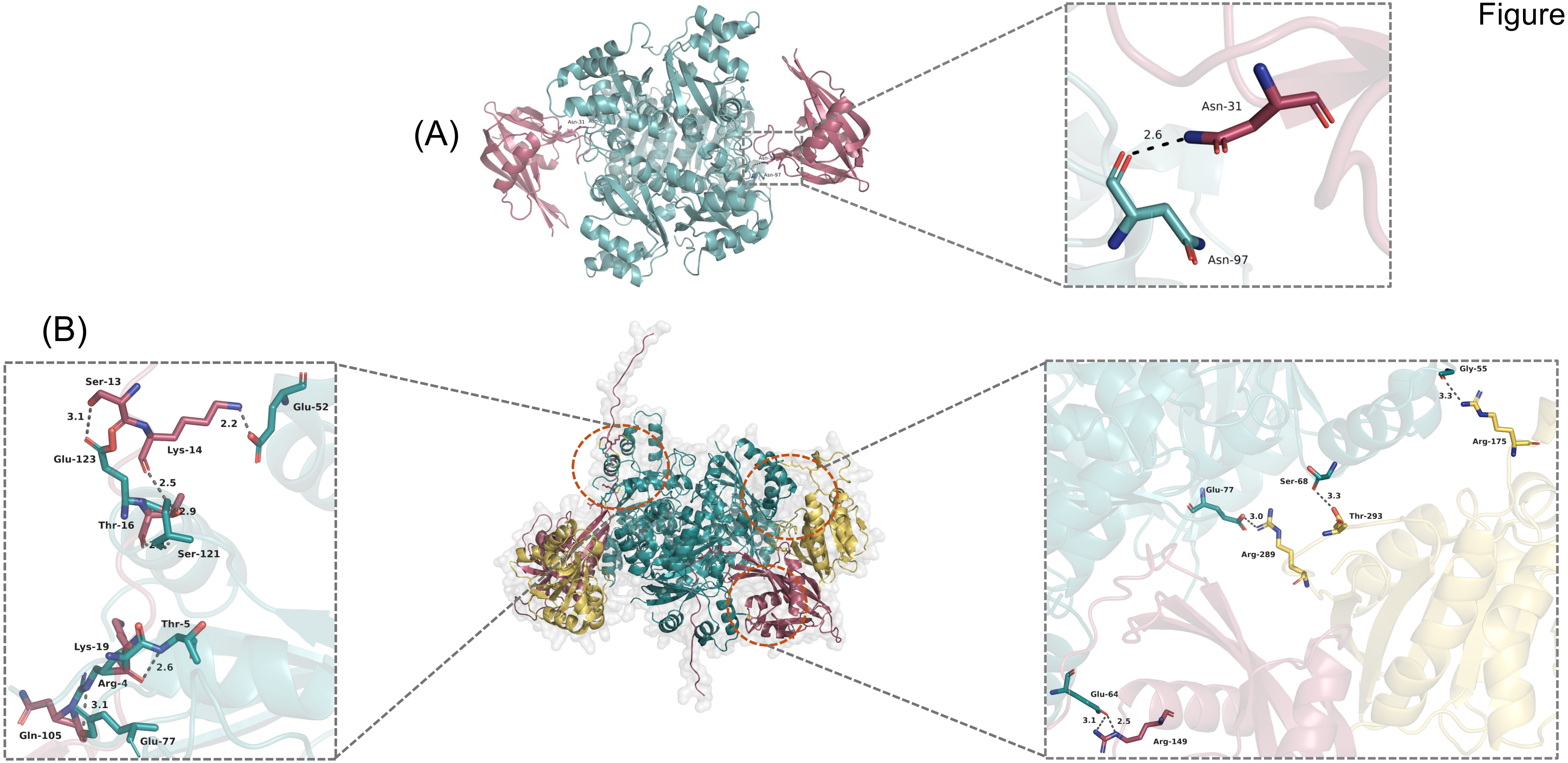


**Figure S6:** Protein-protein interaction prediction using AlphaFold colab. **(A)** The protein sequences of 2X *Eco*MutT (dark red) + 6X *Eco*NDK (cyan) were submitted to AlphaFold colab server for interaction prediction [27]. The results were analyzed using Edu PyMol software [48]. We could detect single interaction between Asn31 from *Eco*MutT (red) with Asn97 from *Eco*NDK (cyan). **(B)**The sequences of 2X *Msm*MutT1 + 6X *Msm*NDK (cyan) were submitted to AlphaFold colab server for interaction prediction [27]. The results were analysed using Edu PyMol software [48]. We could observe interaction of *Msm*NDK with both *Msm*MutT1 domains, *Msm*MutT1 NTD (red) and *Msm*MutT1 CTD (yellow).

### Figure S7

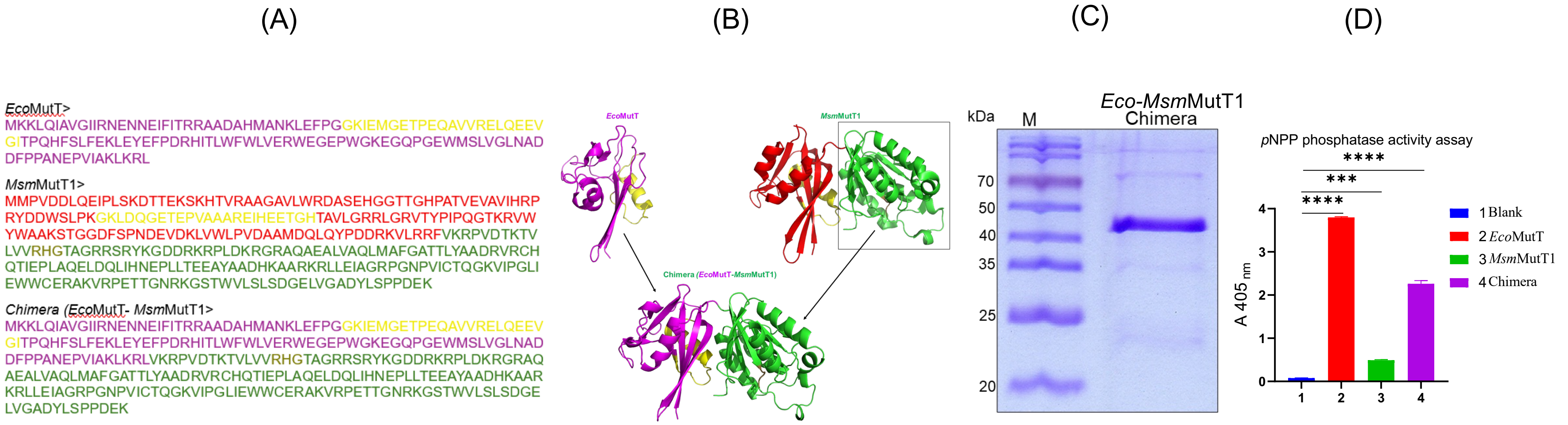


**Figure S7: *Eco*MutT-*Msm*MutT1 (chimera) generation, purification and its phosphatase activity. (A)** Protein sequence of *Eco*MutT (magenta), *Msm*MutT1(NTD red) and (CTD green), and *Eco-Msm*MutT1 (chimera). **(B)** Structure prediction for *Eco-Msm*MutT1 chimera. *Eco*MutT (magenta) PDB entry 1PUN [49], *Msm*MutT1 (NTD red) and (CTD green) (*Ms*MutT1; PDB entry 5GGB [22]) chimera structure prediction [28], [29]. Sequences in yellow represent Nudix hydrolase motifs and in sand colour represent RHG motif. **(C)** Protein purification analysis on 15% SDS-PAGE gel showing the quality of the purified protein. ~3 µg of *Eco-Msm*MutT1 chimera was analysed on 15% SDS-PAGE gel. The calculated molecular mass of *Eco-Msm*MutT1 chimera with $His_6$ tag is ~37 kDa. **(D)** The graph demonstrates the general phosphatase activity of the purified proteins using *p*NPP as a substrate. The substrate was incubated with water (bar 1), 1 µg *Eco*MutT (bar 2), 1 µg *Msm*MutT1 (bar 3) or 1 µg *Eco-Msm*MutT1 chimera (bar 4). Bars represent mean ± SD for n = 3. *p* values, $* p < 0.05$; $** p < 0.01$; $*** p < 0.001$ indicate significant differences between samples; 'ns' represent not significant. One-way ANOVA method was used to calculate *p* value.

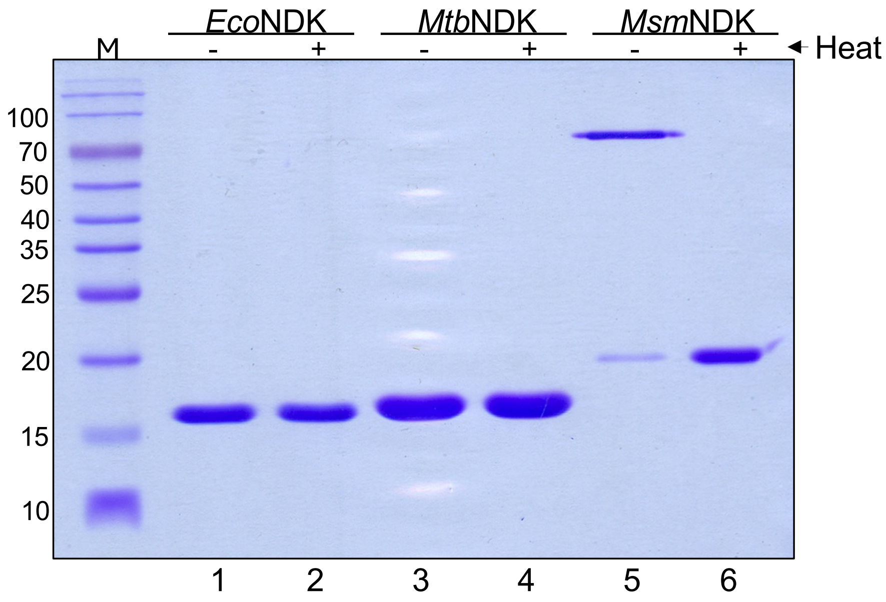


**Figure S8: Effect of heating in SDS-sample buffer on migration of NDK proteins.** Analysis on 12% SDS-PAGE gel showing the effect of heating on the migration of NDK proteins. 1X SDS dye was added to ~2 µg of NDK proteins. Samples were untreated or treated with heat at 90 ºC for 10 min and then loaded onto 12% SDS-PAGE gel. The calculated molecular masses of *Eco*NDK, *Mtb*NDK and *Msm*NDK with His tag are ~16.5, ~16.7 and ~18.1 kDa respectively. Only *Msm*NDK was shown to migrate slowly in unheated condition.

Figure S9

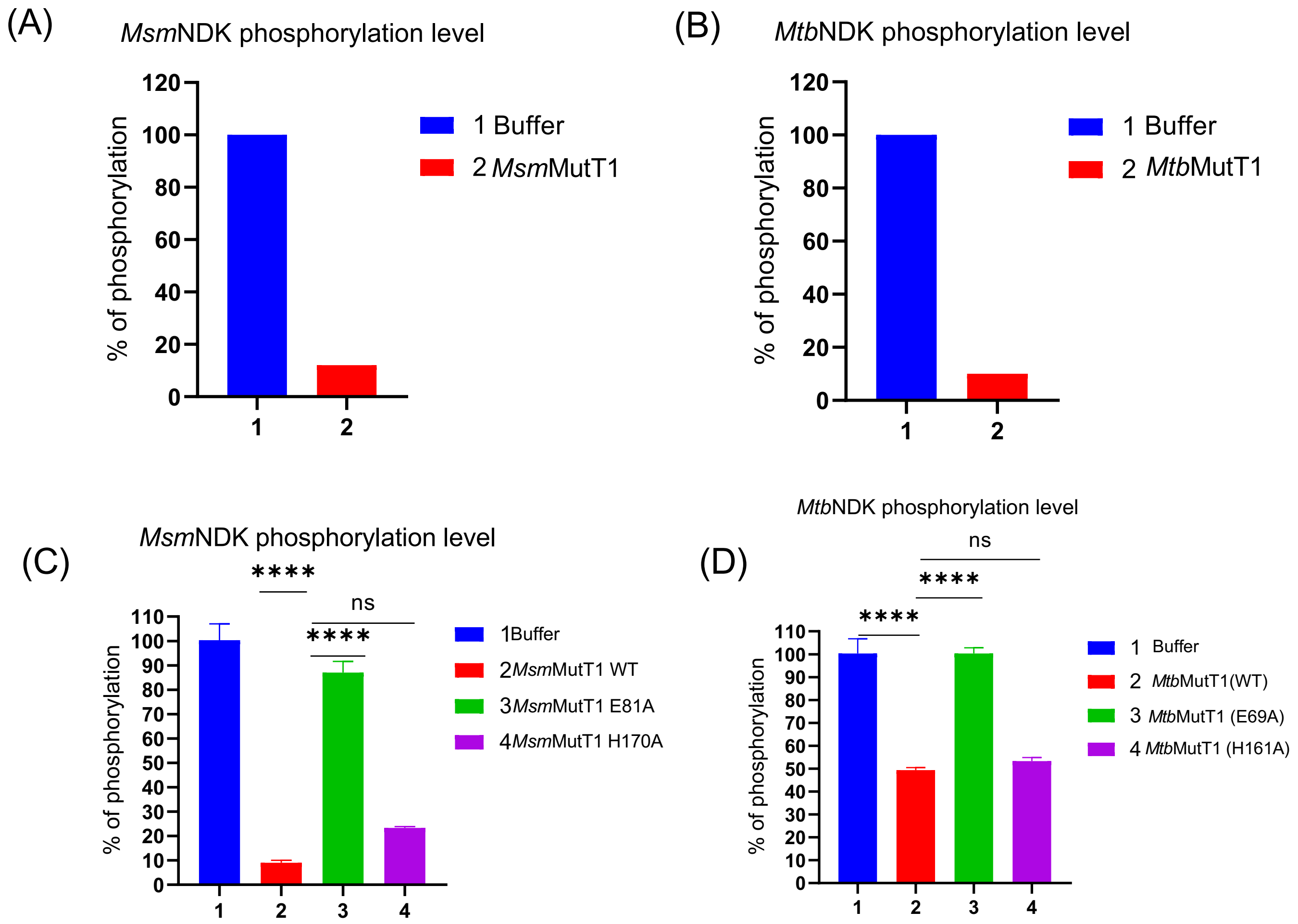


**Figure S9: Quantification of the NDK phosphorylation level. (A)** Phosphorylation levels of *Msm*NDK incubated with buffer or *Msm*MutT1 (bars 1 and 2, respectively). **(B)** Phosphorylation levels of *Mtb*NDK incubated with buffer or *Mtb*MutT1 (bars 1 and 2, respectively. **(C)** Phosphorylation levels of *Msm*NDK incubated with buffer, *Msm*MutT1 *Msm*MutT1 E81A or *Msm*MutT1 H170A (bars 1, 2, 3 and 4, respectively). **(D)** Phosphorylation levels of *Mtb*NDK incubated with buffer, *Mtb*MutT1 *Mtb*MutT1 E69A or *Mtb*MutT1 H161A (bars 1, 2, 3 and 4, respectively). Bars represent mean ± SD for n = 3. *p* values, ** < 0.01; *** < 0.001; ****< 0.0001 indicate significant differences between samples; 'ns' represent not significant. One-way ANOVA method was used to calculate *p* value.

Figure S10

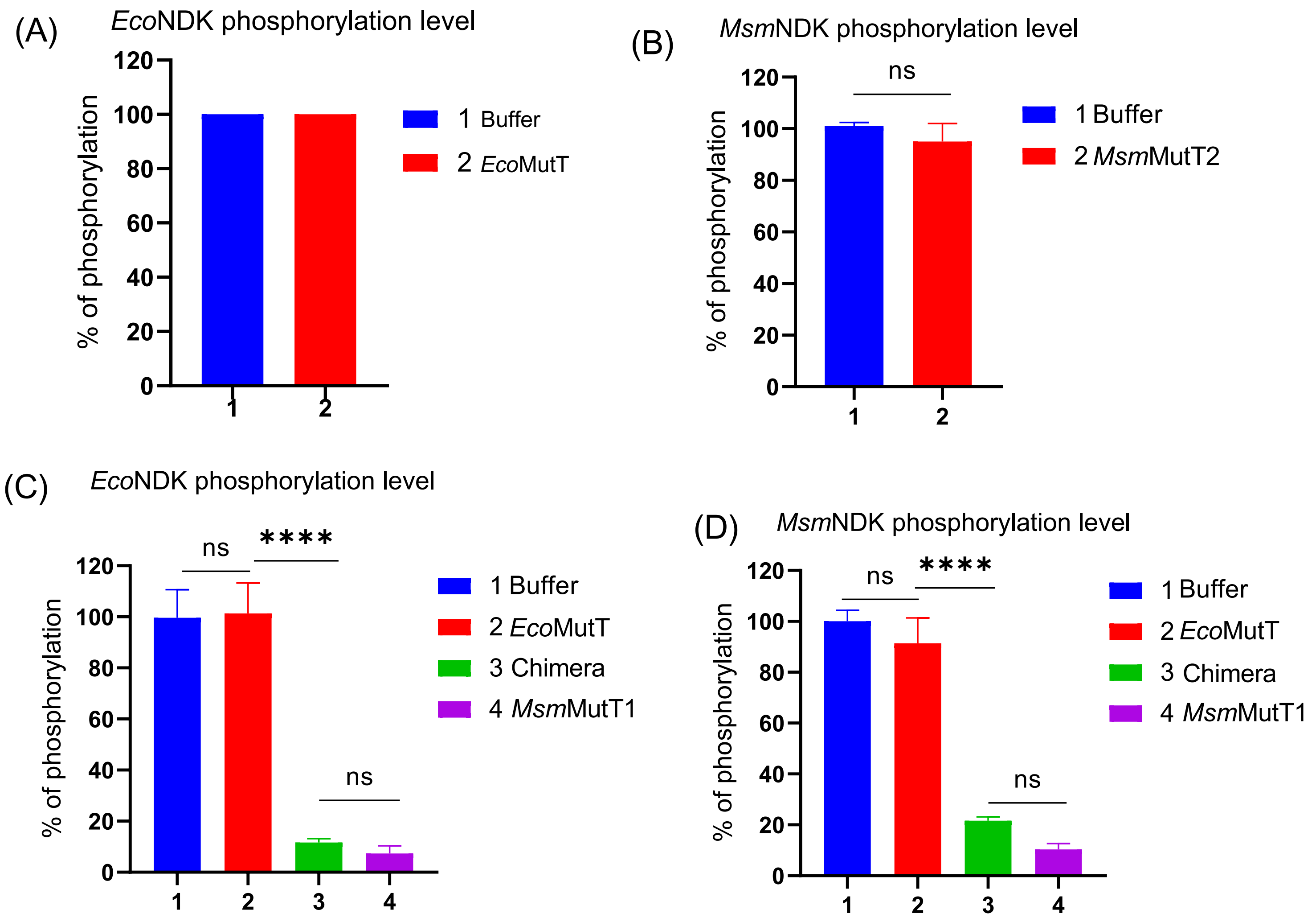


**Figure S10: Quantification of the NDK phosphorylation level. (A)** Phosphorylation level of *Eco*NDK incubated with buffer or *Eco*MutT (bars 1 and 2, respectively). **(B)** Phosphorylation level of *Msm*NDK incubated with buffer or *Msm*MutT2 (bars 1 and 2, respectively). **(C)** Phosphorylation levels of *Eco*NDK incubated with buffer, *Eco*MutT, *Eco-Msm*MutT1 chimera or *Msm*MutT1 (bars 1, 2, 3 and 4, respectively). **(D)** Phosphorylation levels of *Msm*NDK incubated with buffer, *Eco*MutT, *Eco-Msm*MutT1 chimera or *Msm*MutT1 (bars 1, 2, 3 and 4, respectively). Bars represent mean ± SD for n = 3. $p$ values, ** < 0.01; *** < 0.001; **** < 0.0001 indicate significant differences between samples; 'ns' represent not significant. One-way ANOVA method was used to calculate $p$ value.

Figure S11

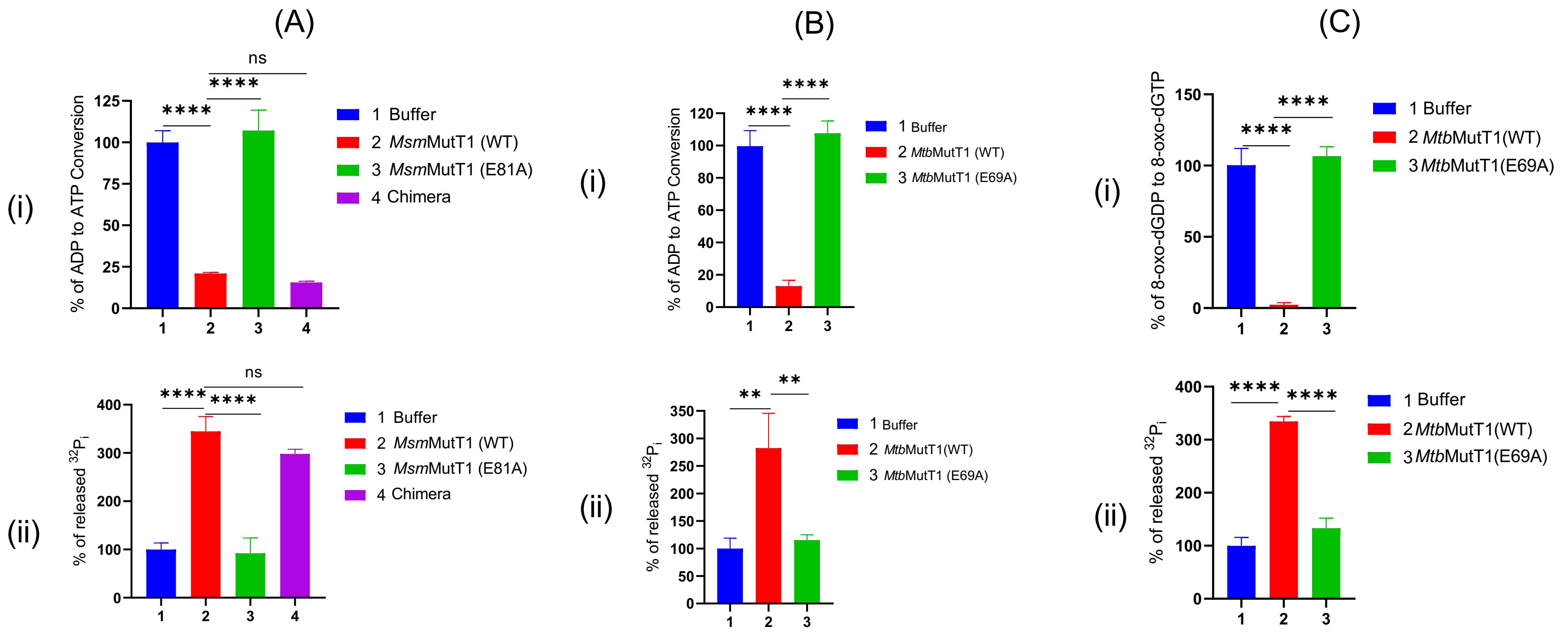


**Figure S11: Quantification of ADP to ATP, and 8-oxo-dGDP to 8-oxo-dGTP conversion by NDK. (A)** (i) % of ADP to ATP conversion by *Msm*NDK in the presence of buffer, *Msm*MutT1(WT), *Msm*MutT1 E81A or *Eco-Msm*MutT1 chimera (bars 1, 2, 3 and 4, respectively) with reference to buffer control (bar 1). (ii) % of $^{32}$Pi release resulting from incubation of *Msm*NDK with buffer, *Msm*MutT1, *Msm*MutT1 E81A or *Eco-Msm*MutT1 chimera (bars 1, 2, 3 and 4, respectively) with reference to buffer control (bar 1). **(B)** (i) % of ADP to ATP conversion by *Mtb*NDK in presence of buffer, *Mtb*MutT1(WT) or *Mtb*MutT1 E69A (bars 1, 2, and 3, respectively) with reference to buffer control (bar 1). (ii) % of $^{32}$Pi released upon incubation of *Mtb*NDK with buffer, *Mtb*MutT1 or *Msm*MutT1 E69A (bars 1, 2 and 3, respectively) with reference to buffer control (bar 1). (C) (i) % of 8-oxo-dGDP to 8-oxo-dGTP conversion by *Mtb*NDK in the presence of buffer, *Mtb*MutT1 or *Mtb*MutT1 E69A (bars 1, 2 and 3, respectively) with reference to buffer control (lane 1). (ii) % $^{32}$Pi release resulting from incubation of *Mtb*NDK with of buffer, *Mtb*MutT1 or *Msm*MutT1 E69A (bars 1, 2 and 3, respectively) with reference to buffer control (lane 1). Bars represent mean ± SD for n = 3. *p* values, ** $< 0.01$; *** $< 0.001$; **** $<$ 0.0001 indicate significant differences between samples; 'ns' represent not significant. One-way ANOVA method was used to calculate *p* value.

Figure S12

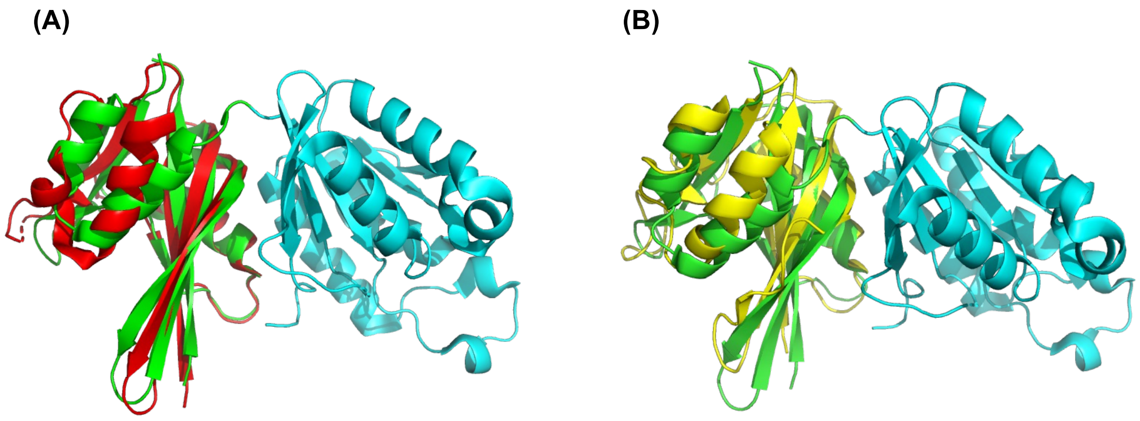


**Figure S12: Structural comparison of *Eco*MutT, *Msm*MutT1, and *Msm*MutT2.** Comparison of *M. smegmatis* MutT1 (*Msm*MutT1 NTD green colour; PDB entry 5GGB; [22]) with its structural homologues, panels: **(A)** *E. coli* MutT (*Eco*MutT red colour; PDB entry 3A6S;[50] ); **(B)** *M. smegmatis* MutT2 (*Msm*MutT2 yellow colour; PDB entry 5ZRG;[51]). The comparison was performed using Edu PyMol software [48].
